# Flies evolved a gyrocompass to maintain navigational accuracy during rapid flight maneuvers

**DOI:** 10.64898/2026.08.18.744964

**Authors:** Ivo G. Ros, Ruoxi Wang, Jaison J. Omoto, William B. Dickson, Michael H. Dickinson

**Author notes:** Correspondence a should be addressed to Michael Dickinson at.

## Abstract

Halteres, the miniaturized hindwings of flies, function as biological gyroscopes that encode angular velocity during flight^1^. Although their established role is to detect flight perturbations and trigger compensatory manoeuvers^2^, here we show that halteres also function as a gyrocompass, maintaining an accurate estimate of heading during rapid turns. To overcome the incompatibility between rapid body rotations and two-photon imaging, we exploited the physics of haltere mechano-transduction to reproduce the inertial forces acting on the halteres during rapid body rotations using micron-scale oscillations of the thorax^3^. Our experiments demonstrate that flies update their internal compass during rapid turns by integrating inertial measurements with predictions derived from efference copy signals. Because body saccades exceed the temporal bandwidth of visual motion processing, this inertial computation allows flies to maintain an accurate internal compass throughout these rapid maneuvers. This capability may enable flies to compute critical parameters such as wind direction and ground speed by comparing measurements made before and after each saccade. Our findings suggest that an ancestral flight-stabilization system was evolutionarily co-opted into a biological gyrocompass, a new function that may have contributed to the adaptive radiation of flies.

---

Dipterans, the true flies, are often acknowledged as the most ecologically diverse order of insects^4,5^. Although the Coleoptera (beetles) and Lepidoptera (moths and butterflies) are both more speciose, flies are noteworthy with respect to their ecological diversity. For example, the Diptera include species whose foraging tactics bear uncanny resemblances to those of dragonflies, moths, bees, wasps, beetles, and fleas^6^, and other species, raising the questions of what specializations allow them to succeed at so many different life histories. One important factor is almost certainly their versatile maggot body form^5^, which enable larval flies to thrive in a diverse array of challenging habitats. As adults, however, the feature that sets flies apart from nearly all other insects is their possession of halteres—the highly modified hindwings that function as equilibrium organs during flight^1,7,8^.

Each haltere consists of a spherical knob attached to the body via a narrow stalk equipped with ∼150 strain-sensitive sensitive campaniform sensilla (CS) that project into the ventral nerve cord (VNC)^1,9,10^ (Fig 1a,b). Evidence suggests that the halteres function both as a clock and a gyroscope^11^. The clock function arises through the detection of the large inertial forces generated as the halteres accelerate back and forth, which provide timing signals to ensure that wing control muscles fire at biomechanically effective phases of the stroke cycle^12,13^. The gyroscopic role depends on a subset of CS sensitive to the much smaller Coriolis forces that deflect the halteres from their stroke plane when the body rotates^1,3^. Because the time history of Coriolis forces depends on the orientation of the body’s angular velocity vector, flies can use their halteres to discriminate angular velocity about all three rotational axes (yaw, pitch, and roll)^14,15^. Whereas the role of halteres in equilibrium reflexes is well established^2^, we test here the hypothesis that halteres also function as a gyrocompass via their input to the Central Complex (CX), a brain region specialized for navigation and vector computation^16^ (Fig. 1).

**Fig. 1.**
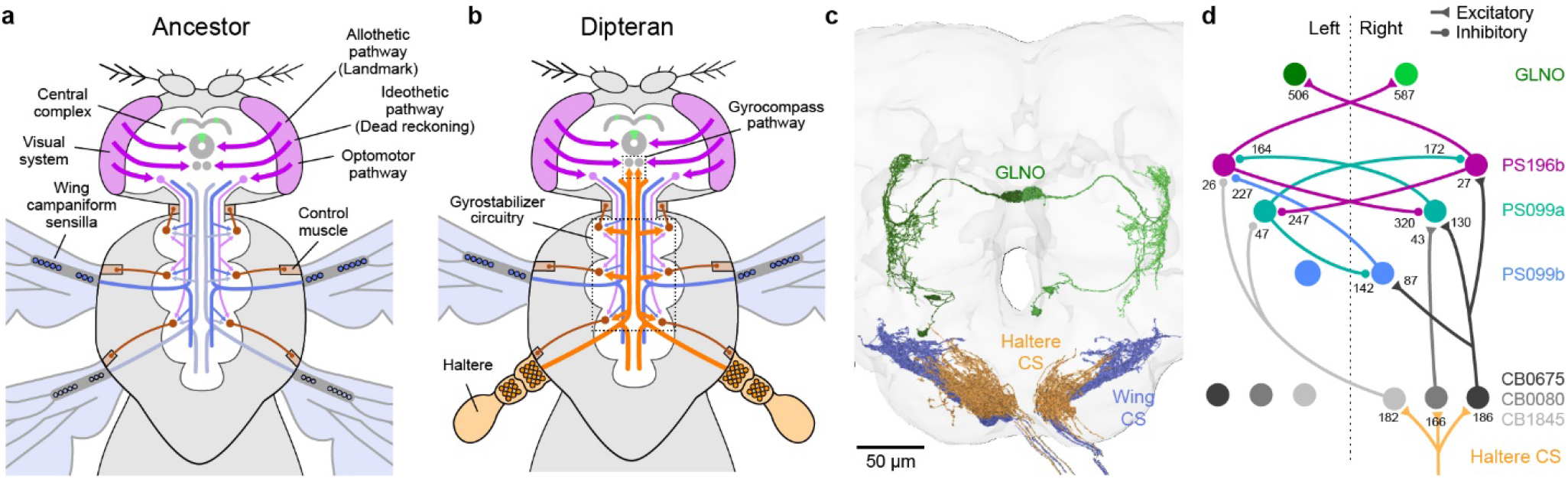
Gyrocompass hypothesis. **a**, The ancestor of all Diptera possessed two sets of wings, each equipped with campaniform sensilla (CS) and control muscles. Flight control was mediated by a combination of local reflexes and descending pathways, while the compass within the Central Complex (CX) was updated by a combination of idiothetic and allothetic visual cues. **b**, In the course of evolution, the hindwings transformed into halteres replete with specialized sensilla that encode Coriolis forces and thus angular velocity. Together with specialized circuits within the VNC, the halteres constitute a gyrostabilizer that maintains stability in response to perturbations. We propose that the halteres also play a role in navigation akin to the gyrocompasses of ships and aircraft, mediated by novel connections to compass circuitry in the CX. **c**, Projections of ascending wing and haltere CS **neurons** in the subesophageal zone along with GLNO neurons in the CX. **d**, Ascending pathway linking ipsilateral projection of haltere afferents to the GLNO cells. Numbers indicate synapse counts; see Methods for details of the analysis. Neuroanatomy in panels c and d is based on the FlyWire-FAFB connectome^17,18^.

## Micron-scale oscillations of the thorax simulate body rotation

Testing the gyrocompass hypothesis is challenging due to the difficulty of rapidly rotating a fly while recording from small neurons in its brain. To circumvent this hurdle, we adopted an ingenious approach developed by Gerbera Nalbach for investigating gaze stabilization in blowflies^3,14^. Nalbach showed it is possible to accurately replicate Coriolis forces by subjecting a fly’s body to small-scale linear oscillations (See Methods and Supplemental Information). Using modern electronics and fabrication techniques, we miniaturized her approach for use with *Drosophila* (Fig. 2). The key components of our apparatus, nick-named the *Flybratron*, are shown in Fig. 2a&b. We attach a fly to a voice coil oscillator via a tungsten pin, and use an optical wingbeat analyzer to track the shadows of its wings cast by an infrared LED^19^, thus generating a stable synch pulse near the start of each downstroke. Clocked by this trigger, a microcontroller calculates the acceleration required to simulate the Coriolis forces. To simulate yaw (Fig. 2a-f), the necessary waveform of the thorax oscillations is a harmonic function at twice the wingbeat frequency, a pattern that forces each haltere into a narrow figure-of-eight as they beat up and down.

**Fig. 2.**
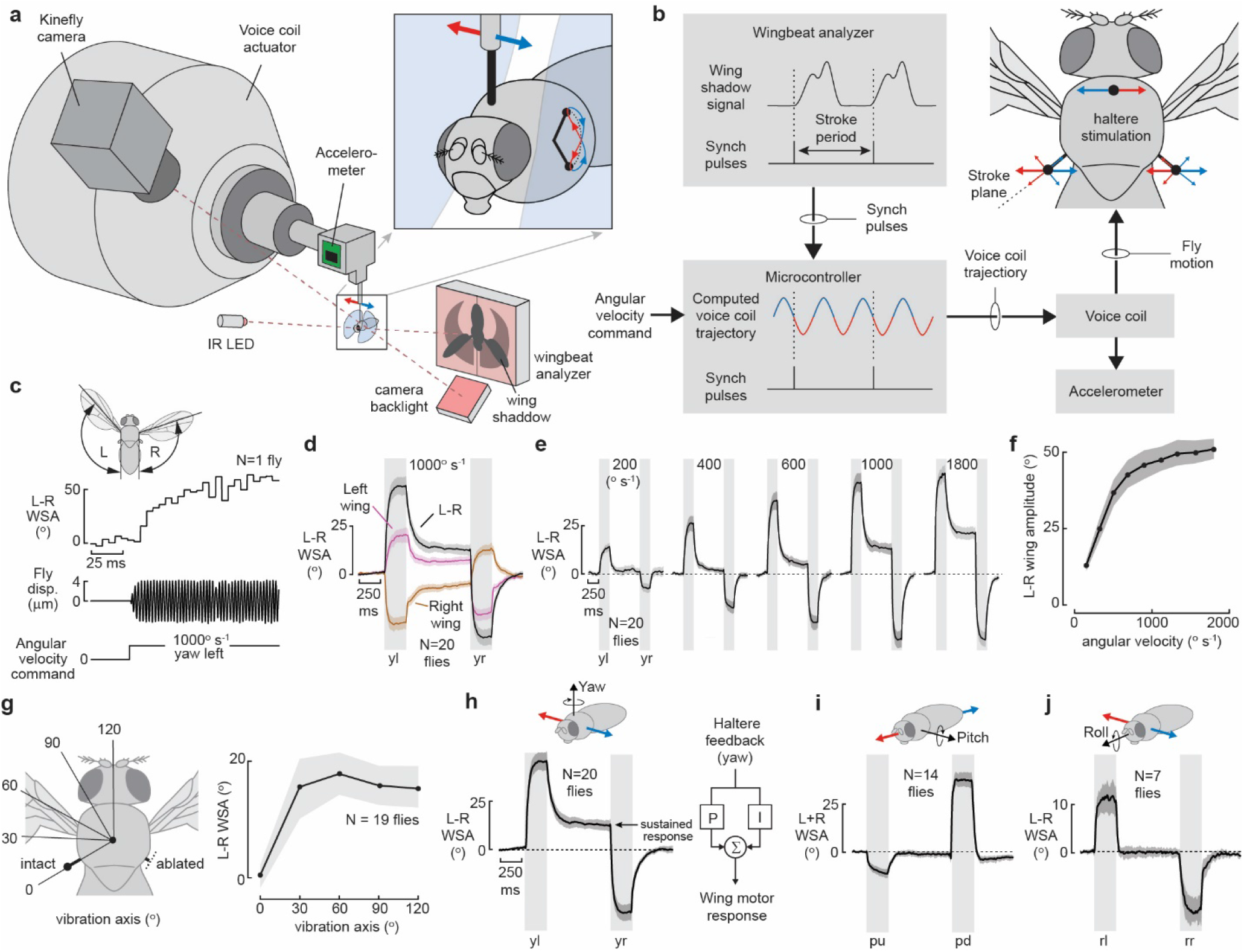
Micro-oscillation accurately simulates Coriolis forces on the halteres. **a**, Experimental setup. A voice coil oscillator vibrates a tethered fly while its wing motion is tracked by a wingbeat analyzer and machine vision system *(Kinefly)*. **b**, Schematic view of the Flybratron. The wingbeat analyzer detects a synch pulse each wingbeat for creating the phase-locked acceleration of the fly’s body that simulates the Coriolis forces acting on the halteres during body rotation. **c**, Example data of a compensatory changes in wing motion induced by a simulated leftward yaw rotation at 1000^°^ s^-1^. **d**, Compensatory response to yaw left (yl) and yaw right (yr) stimuli. Here and throughout, data are plotted as baseline-subtracted mean and 95% confidence intervals (CI) with the sample size indicated in the panel. **e**, Response to a series of left and right yaw rotations at different angular velocities. **f**, Angular velocity response curve. **g**, Ablation experiments designed to test sensitivity of the halteres to axial loading. Flies with one haltere ablated respond to simulated rotation, provided the vibration axis is not parallel to the stroke plane of the intact haltere (0°) **h**, Data from panel d replotted to emphasize the sustained response following the first yaw rotation, consistent with the presence of Proportional-Integral (PI) feedback (inset at right). **i, j**, Responses to simulated pitch and roll are robust and stereotyped but do not show evidence of an integral term (pu and pd indicate pitch up and down; rl and rr indicate roll left and right). See also Supplementary Video 1.

The latency of compensatory reflexes evoked by simulated rotation is remarkably short; we detected changes in wing motion within 1.5 wingbeats (∼7 ms) following the onset of a yaw stimulus (Fig. 2c). Due to scaling effects, Nalbach’s method developed for a much larger fly is particularly effective in *Drosophila*—an oscillation amplitude of 4 μm replicates an angular velocity of 1000 ° s^-1^ (see **Supplementary Video 1**, Supplementary Information). The reflexes were stereotyped and reproducible across both trials and animals, highlighting the hard-wired nature of the relevant circuits within the VNC^20^. Behavioral responses, which we measured as the bilateral difference in wingstroke amplitude (L-R WSA), rose monotonically to 50^°^ (Fig. 2e-f), close to the biomechanical limit for the range of wing motion^21,22^. Consistent with prior studies employing actual body rotation^15,23^, flies responded to simulated yaw rotation via increases in the stroke amplitude of one wing and decreases in the other (Fig. 2d).

To gain initial insight into how flies coordinate bilateral sensory signals within the VNC, we measured the responses to yaw rotation after removing one of the flies’ two halteres. The measured motor responses of both wings were roughly 50% of that measured in intact flies, confirming that the information for one haltere is sufficient to detect yaw, and that flies appear to sum the yaw signals encoded by its two halteres^15,24^. Flies exhibited no compensatory reactions if oscillated parallel to the long axis of the intact haltere (Fig. 2g), verifying that the CS encoding Coriolis forces are insensitive to axial loading^1,24^.

By adjusting the orientation and frequency of the Flybratron, we could also simulate rotation about the pitch and roll axes (see Methods, Supplementary Information). Again, the wing motor responses to simulated rotation were consistent with prior studies using actual rotation (Fig. 2i, j)^15,25^. Flies responded to yaw with a rapid change in wing motion followed by a sustained response that continued long after the stimulus ended. This sustained response immediately ends, however, when the fly is subjected to the same rotation in the opposite direction (Fig. 2h). These results support a hypothesis, based on free flight perturbation experiments, that flies implement Proportional-Integral (PI) control in their compensatory reflexes to yaw rotation^26^ (Fig. 2h, inset). Feedback control with an integral term would be beneficial for correcting small rotational drift about the yaw axis during flight, thus enabling the animals to fly much straighter than they could otherwise^27^. We found no evidence, however, for a sustained component suggestive of PI control in the responses to either roll or pitch (Fig. 2j, k)^28–30^.

### The halteres provide angular velocity information to compass neurons

To test if signals from the halteres are used by the compass network, we modified the Flybratron so it was compatible with 2-photon imaging (Fig. 3a, b). Whereas the fly’s head was attached to a custom-built physiology stage as in prior studies^31,32^, we glued the fly’s thorax to a horizontal tungsten wire extending from the voice coil. The movement artifacts caused by oscillating the fly were comparable to what we typically observe in these preparations due to the flapping movements of the wings, and were easily corrected by an image-registration protocol (see Methods).

**Fig. 3.**
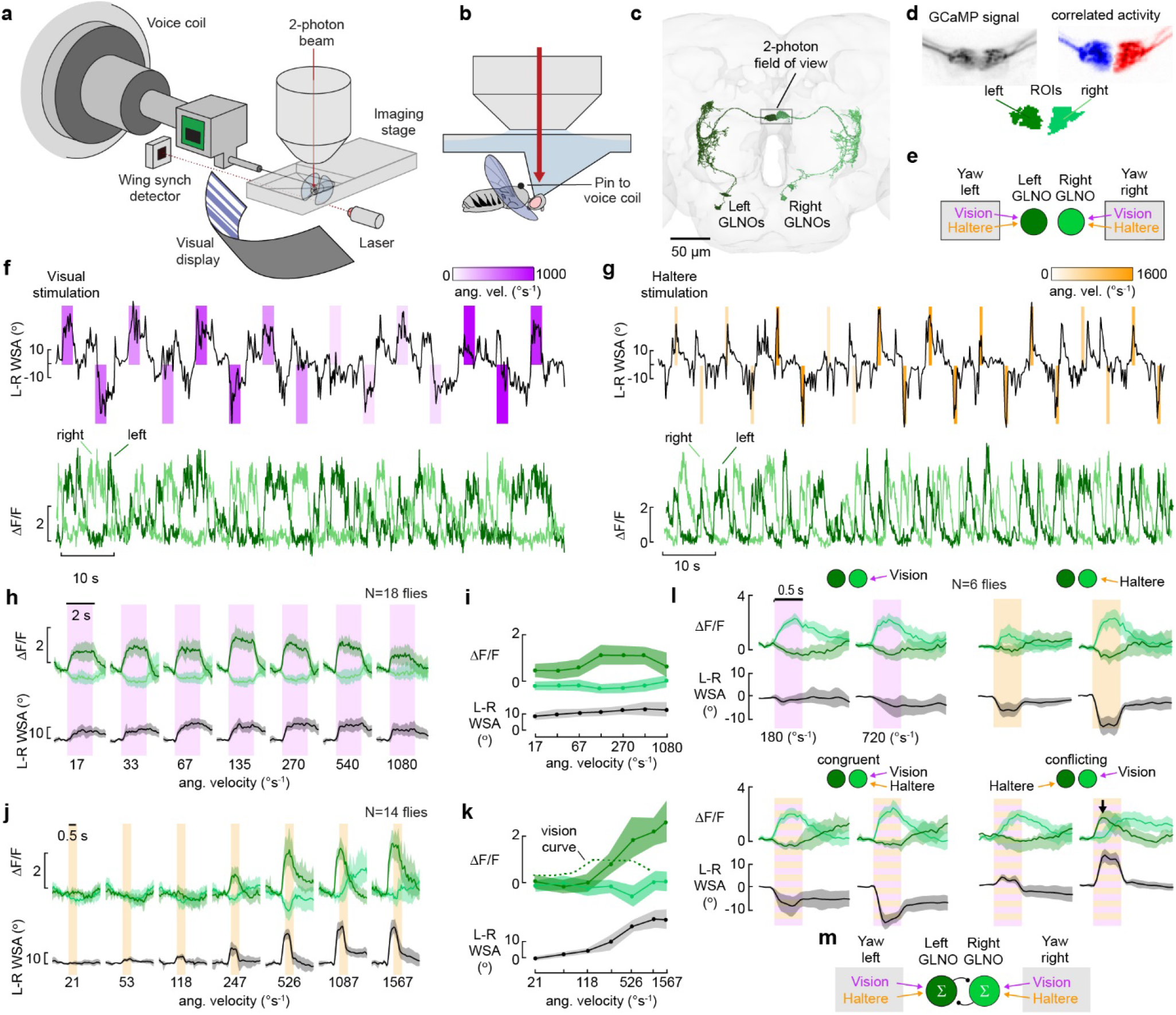
Angular velocity estimates from halteres and visual system converge on GLNO neurons. **a**, Modification of Flybratron for functional imaging. **b**, The voice coil oscillates the thorax side-to-side via a pin attached to the notum, while the fly’s head is fixed to the imaging stage. **c**, Two-photon field of view superimposed on images from FlyWire-FAFB^17,18^. **d**, Signals from left and right GLNO neurons are segmented according to correlated activity. **e**, Schematic of how vision and haltere input converges on GLNOs. **f**, Example traces showing responses to visually simulated yaw. The top trace (black) shows the steering responses of the fly superimposed with purple patches encoding stimulus velocity and direction. The bottom traces (light and dark green), show the activity of left and right GLNOs. **g**, similar to f, but for haltere stimulation (orange patches). **h**, GLNO responses to visual rotation parsed according to stimulus velocity. **i**, Data from h plotted as a function of angular velocity; note that the abscissa is a geometric scale. **j-k**, Similar to h and i, but for haltere stimulation. In panels h-k, data in response to yaw right were reflected so they could be combined with responses evoked by yaw left. **l**, GLNO activity in response to simultaneous input from visual system and halteres. The top panels show responses to vision-only and haltere-only input at low and high angular velocities. The bottom panels show congruent and conflicting input. At the start of the conflicting stimuli at 720 ° s^-1^, the left GLNO, which transiently shows elevated activity (black arrow). **m**, The results of the conflict experiments support a simple model in which the two modalities sum linearly on the GLNOs, with the two bilateral circuits organized as a winner-take-all motif, here schematized as reciprocal inhibition. See also Supplementary Video 2.

Prior work identified the GLNO neurons as critical nodes in transmitting sensory and motor signals encoding body rotation to compass neurons in the Central Complex^33^. The pair of nearly identical cells (GLNOi,ii) receive input from the Gall and Lateral Accessory Lobe on each side of the brain and make synapses with PEN neurons in the noduli that project to the contralateral Protocerebral Bridge (PB)^34^. We determined that these cells are glutamatergic and thus inhibitory (Extended Data Fig. 1). As a result of this circuitry, idiothetic signals encoding clockwise rotation of the fly cause a counterclockwise movement of the EPG activity bump and vice versa^35,36^. Prior measurements in non-flying flies show that the GLNO neurons are broadly tuned to the angular rotation of large-field visual patterns rotating about the yaw axis, but this response attenuates at speeds above 100 ° s^-1^, well below the angular velocities that flies experience during fast flight maneuvers^33,37^. Using the FlyWire connectome^18^, we identified a 3-cell pathway consisting of the CB0675, PS099(a&b), and PS196(b) cells that links haltere CS afferents in the Subesophageal Zone to the GLNO neurons (Fig. 1b, c). This anatomical pathway in *Drosophila* is consistent with extracellular recordings in quiescent flesh flies demonstrating that haltere movements elicit spikes at locations throughout the CX^38^. We performed an identical connectomic analysis on the serially homologous population of campaniform neurons at the base of the wing, but found no evidence for a functional pathway to the GLNO cells. Thus, the pathway linking haltere information to the CX has been modified from the ancestral condition when the hindwing still played an aerodynamic role (Fig. 1a, b).

To test the predictions from the connectome, we recorded the activity of the left and right GLNOs by expressing the Ca^+2^ indicator, GCaMP7f, under control of a driver line (BDSC_39933) that targets both GLNO cells (Fig. 3d). While imaging from terminals in the noduli, we subjected flies to virtual rotation across a range of angular velocities via either wide-field visual motion (Fig. 3f, h, i) or haltere stimulation (Fig. 3g, j, k, **Supplementary Video 2**). Because the dynamics of motion vision are slower, we presented the visual stimuli for 2 s, compared to 0.5 s for haltere stimuli. The polarity of the responses were consistent with previous recordings from non-flying flies^33^; rightward visual motion simulating a yaw rotation to the left elicited an increase in the activity of the left GLNO and a decrease in activity of the right GLNO, with leftward visual motion having the opposite effect. As in prior measurements in walking flies^33^, the visual responses of GLNO neurons during flight exhibited band-pass characteristics with a broad peak near 100 ° s^-1^ (Fig. 3i). In contrast, GLNO responses mediated by haltere stimulation exhibited high-pass characteristics, with the neuronal responses rising steeply at higher angular velocities (Fig. 3k).

Throughout each trial, the responses exhibited a ‘flip-flop’ pattern of activity in which the right and left pairs of GLNOs were either on or off, but rarely active at the same time (Fig. 3f, g). The flip-flop nature of the circuitry was especially noticeable when we presented conflicting stimuli such as rightward rotation at 720 ° s^-1^ via the halteres paired with conflicting leftward rotation via the visual system (Fig. 3l, lower left panel). Initially the left GLNO activity was high while the right GLNO activity was low, but this polarity reversed after ∼0.5 s, presumably because as the slower visual input rose, it eventually exceeded the contralateral haltere signal and flipped the bilateral pattern of GLNO activity. These experiments support a prior work indicating that the bilateral GLNO pathways are wired to generate a “winner-take-all” outcome^33^.

### Gyroscopic feedback from the halteres moves the EPG activity bump

Our results indicate that sensory information from the halteres reaches the GLNO neurons, but they do not provide quantitative measurements for how far the compass bump rotates in response to a rotational stimulus. To do this, we recorded the population activity of EPG neurons in the PB and again presented flies with rotational stimuli over a range of different angular velocities (Fig. 4, **Supplementary Video 3**). In these experiments, we chose the same stimulus duration (0.5 s) for both visual and haltere stimuli so that the absolute angular rotation of each stimulus would be the same for the two modalities. The results were consistent with the GLNO recordings (Fig. 4) in that the magnitude of bump rotation was relatively flat with respect to angular velocity for the visual stimulus (Fig. 4c) but rose steeply with the angular velocity signal delivered via the halteres (Fig. 4e).

**Fig. 4.**
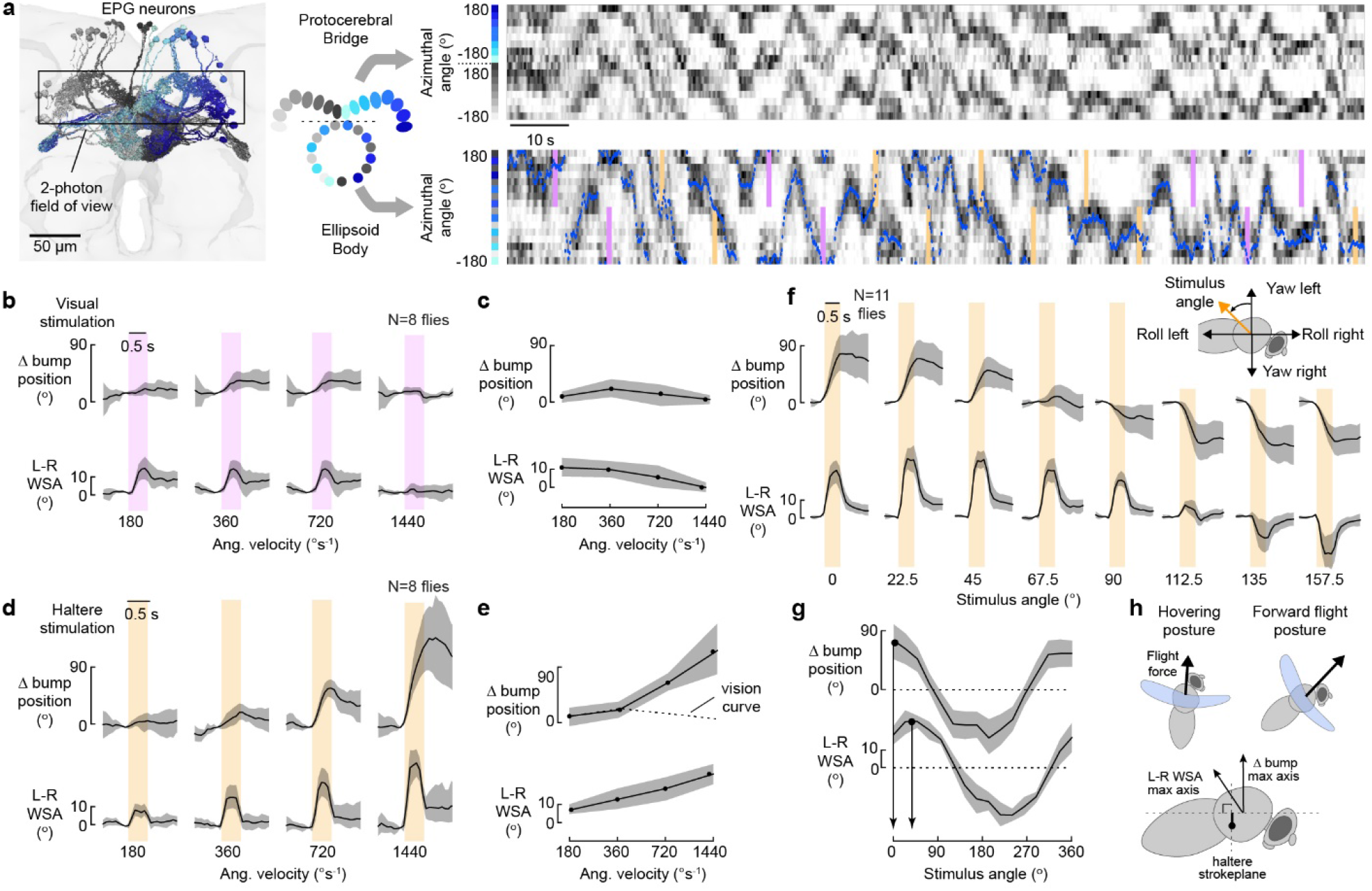
The compass heading in the EPG network rotates in response to haltere input. **a**, EPG network and the field of view used in 2-photon experiments. As in prior studies, data from the left and right halves of the PB were remapped to create a single azimuthal representation of the heading bump (dotted blue trace). Flies were presented with a randomized set of visual (purple) and haltere (orange) stimuli at four different angular velocities. **b-c**, EPG bump movement and steering behavior in response to visual rotation at four different angular velocities. **d-e**, same as **b** and **c**, but for haltere stimulation. **f-g**, EPG bump movement and steering behavior in response to the direction of simulated angular velocity in the roll-yaw plane. **h**, Although flies adopt different postures at different flight speeds, the input from the halteres to the compass system appears fixed to a rotation axis oriented parallel to the halter stroke plane, which is roughly perpendicular to the longitudinal body axis. In contrast, the L-R WSA response is maximal about a rotation axis that is inclined ∼40^°^ caudally.

### The halteres report angular velocity relative to the longitudinal body axis

For a walking insect, the longitudinal body axis is roughly parallel to the 2-dimensional plane in which it moves; thus, it would make sense for the compass system to report heading with respect to this orientation. When flying, however, flies actively adjust their posture with respect to the horizontal plane—adjustments that alter the horizontal component of the mean aerodynamic force vector as necessary for different flight speeds^39,40^ (Fig. 4h). As a consequence, the axis around which flies change heading in flight is not necessarily aligned with their longitudinal body axis. To determine around which body axis the halteres report angular velocity to the compass during flight, we programmed the Flybratron to oscillate with waveforms that were a linear combination of signals simulating roll and yaw, thus allowing us to present rotational stimuli around any arbitrary axis in the yaw-roll plane (Fig 4f). The resulting tuning curve (Fig. 4g) shows that the compass system is most sensitive to rotations about flies’ longitudinal body axis. This result is consistent with the expected magnitude of the Coriolis forces, which are greatest when the rotational axis of the body is parallel to the stroke plane of the halteres. In contrast, we found that the magnitude of the wing steering response (L-R WSA, Fig. 4g) was tuned to an axis tilted by ∼40^°^ rostral to the longitudinal body axis, which is consistent with a linear sum of the separate wing motor responses to roll and yaw as previously reported^15,23^. Thus, the mapping of body rotation to bump motion is similar in walking and flying flies, even though body orientation varies more in flight due to the changes in body orientation with air speed. One means by which flies might deal with the changes in body posture during flight would be to actively maintain their head at a constant pitch angle relative to the ground^41,42^.

### Flies update their compass during saccades by a combination of prediction and measurement

Like many other fly species, *Drosophila* exhibit rapid turns called saccades in which they change heading by ∼90^°^ in ∼100 ms^37,43^. Flies use directed saccades to avoid looming threats and obstacles, but also generate them spontaneously at a rate of ∼0.5 Hz in the absence of sensory stimuli^44^. Spontaneous saccades are much more kinematically stereotyped than collision-avoidance saccades^45^, suggesting that the animals may be attempting to change course by the same amount each time. The behavioral role of saccades is not well understood, but it has recently been suggested that they might serve a role in active sensing during flight^46–49^. Prior studies of flies walking on a spherical treadmill show that fictive turns are accompanied by a rotation of the heading bump even when visual cues are absent^35,36,50^, and we see a similar phenomenon in flying flies (Fig. 5a). The efference copy responsible for this bump movement likely converges on the GLNO cells^33^. To investigate if and how this predictive input to the compass system is integrated with haltere feedback during the rapid turns, we programmed an online filter to detect the execution of spontaneous saccades and then immediately trigger a haltere stimulus that coarsely mimics the rapid rotation a fly would experience during a free flight saccade (Fig. 5c). In the absence of haltere feedback, we measured bump rotations of ∼40^°^ when flies executed a spontaneous saccade. By comparison, the bump rotated by ∼50^°^ when presented with a haltere stimulus that simulated a 90^°^ turn. When the efference copy signal and re-afferent feedback from the halteres were both present, flies exhibited a bump rotation of ∼90° —a value that is remarkably close to the measured magnitude of free flight saccades^43,45^. These results suggest that compass rotation during a saccade uses both predictive information from the motor system and sensory measurements from the halteres. Prior authors have noted that the means by which flies estimate bump position using sensory feedback resembles a circular Kalman filter^51^. Our experimental data from flying flies supports and extends this idea; the process of updating compass heading using efference copy and haltere feedback is analogous to a Kalman filter with fixed gains for the predictive and measurement terms. Presumably, flies benefit from this approach because the noise associated with the prediction and measurements are independent, and thus they achieve a better estimate of the true saccade magnitude using a weighted average. Motor noise is particularly problematic for saccades because these rapid maneuvers are controlled by very few motor units; the addition of just one additional motor neuron spike in some phasic steering muscles would have substantial consequences on the resulting pattern of wing kinematics^52,53^.

**Fig. 5.**
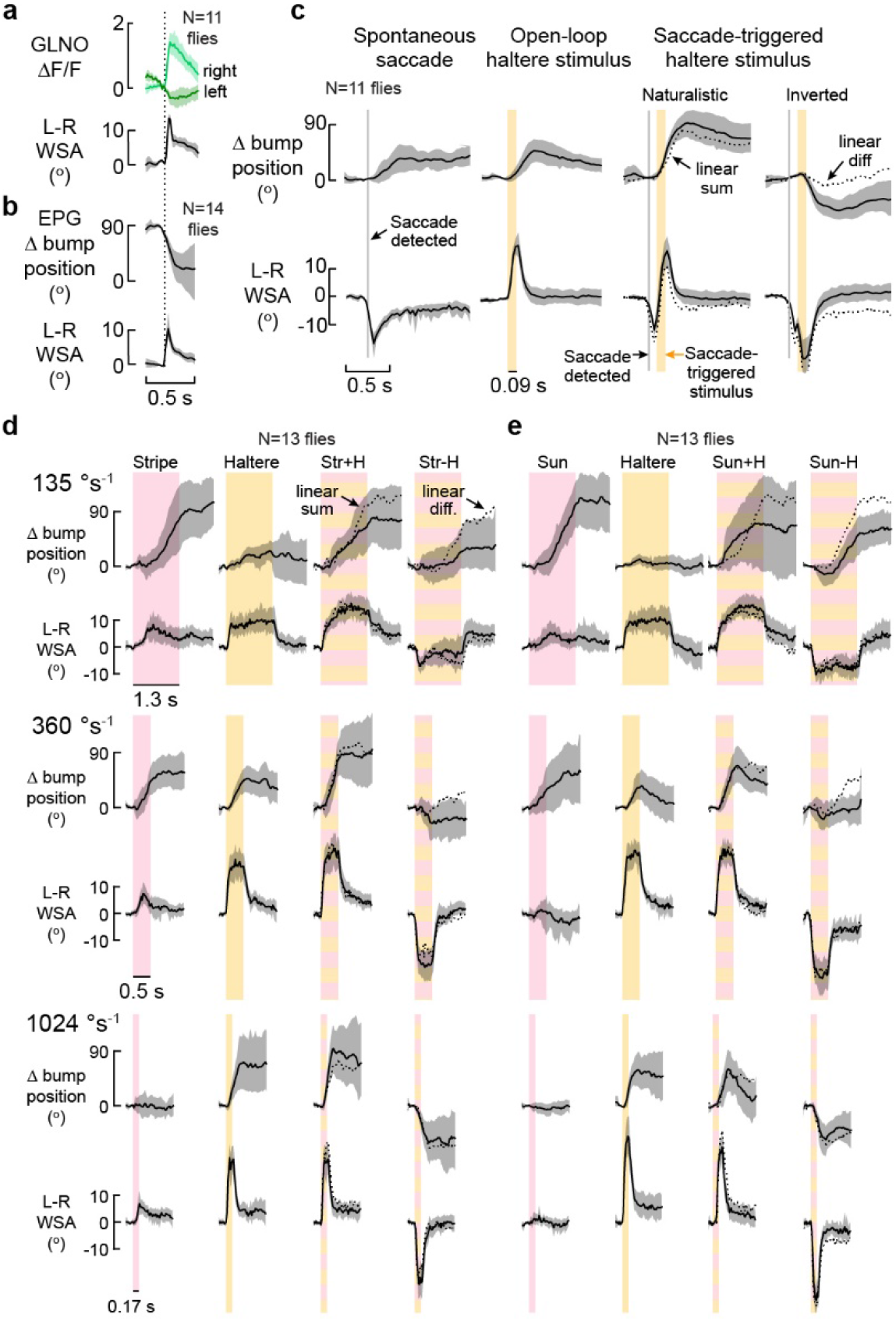
During flight saccades, the compass rotates accurately via a combination of haltere measurement and motor prediction. **a**, GLNO activity during spontaneous saccades. **b**, EGP bump position during spontaneous saccades. In a and b, saccades to the left have been reflected and combined with saccades to the right. **c**, EPG bump position and wing motion in response to simultaneous input from efference copy signals and halteres. Left-most panel shows changes in wing motion and bump position during saccades, detected by a real-time trigger. Again, saccades to the right have been reflected and combined with saccades to the left. The next panel to shows response to 90 ms haltere stimuli presented at 1000 ° s^-1^. In the two panels on the right, haltere stimulation with either a naturalistic or inverted polarity was presented immediately after a spontaneous saccade was detected; linear predictions are shown by dotted lines. **d**, EPG bump position and L-R WSA steering behavior in response to rotation of a bright, rotating stripe and/or a haltere stimulus at angular velocities of 135, 360, and 1040 ° s^-1^. The responses to the rotational landmark and haltere only are shown in the first two columns; the next two show naturalistic (Str+H) and conflicting (Str-H) cases of simultaneously presented stimuli. **e**, same as **d**, but with an ersatz sun stimulus as the landmark.

### Navigation via a combination of dead-reckoning and landmark orientation

Our results suggest that flies adjust the position of their compass orientation during rapid turns using a combination of idiothetic cues from the visual system and halteres, combined with a prediction based on efference copy. Collectively, this constitutes a navigational system that operates via dead reckoning, i.e. estimating changes in heading by integrating angular velocity. In addition, flies possess a landmark-based system that links the compass to the external world via a large population of ring neurons that synapse directly on EPG cells in the Ellipsoid Body^34,54^. Each population of ring neurons consists of cells that are selective to a particular external cue (e.g. the position of the sun, or pattern of skylight polarization) with receptive fields that tile azimuthal space^55^. The map of these external cues onto the EPG cells is not hard-wired, but rather is constructed via a Hebbian learning rule^56–58^, thus providing a means of flexibly yoking the EPG bump to whatever landmarks are currently available. In principle, landmark–based navigation is more accurate over long distances because the integration of angular velocity upon which dead reckoning depends is suspect to drift. However, ships, airplanes, and animals all face circumstances when landmark cues are obscured, during which time dead reckoning provides a viable alternative.

To investigate the functional convergence of the ring neurons and GLNOs, we presented flies with rotating landmarks (bright vertical stripe or ersatz sun) across a range of angular velocities and compared the resulting EPG bump movement with the same rotation delivered via the halteres (Fig. 5d, e). We set the stimulus duration so that the simulated rotation was 180^°^ for each stimulus, regardless of the angular velocity. The results were consistent across experiments and independent of which landmark cue we presented. At 135 ° s^-1^, the landmark spun the bump by ∼100°, but its impact diminished with increasing rotational speed. At 1040 ° s^-1^, the bump could no longer follow the rotating cue. In contrast, the influence of the halteres increased with stimulus speed, with the strongest response at 1040 ° s^-1^. When we presented both stimuli simultaneously, the net effect of the motion of the bump indicated a roughly linear sum of the two independent responses. Prior closed-loop experiments on flying^59^ and walking^50,60^ flies demonstrate that the EPG bump moves appropriately when a bright vertical bar is instantaneously ‘jumped’ from one azimuthal position to another, which would seem inconsistent with our measurements showing no bump movement in response to a 1040 ° s^-1^ rotation of a landmark. This discrepancy may arise because bump rotation following a landmark jump takes several seconds to complete, much longer than the time course of our fast rotation experiments. Our results suggest that the ring neuron system is not well suited for tracking rotation of a landmark during a rapid rotation and thus cannot provide an accurate measure of heading immediately following the completion of a fast manoeuvre. In contrast, the combination of haltere feedback and a prediction based on efference copy would provide a reasonable estimate of the change in heading immediately after executing a saccade (Fig. 5c).

## Discussion

These results expand our understanding of the unique functional advantages of dipteran halteres; not only do flies use their unique gyroscopic sensors for stabilization, they exploit them to increase the precision of their internal compass (Fig. 6a). Vision- and haltere-based measurements of angular velocity sum at the level of the GLNO neurons (Fig. 3), a convergence that greatly extends the range of angular velocity that the navigation system can integrate. The GLNOs also receive an efference copy signal during fast turns, constituting a simple Kalman filter in which the compass heading during rapid turns is updated by a weighted sum of prediction and measurement (Fig. 6a). Although the GLNOs and ring neurons rotate the EPG bump via quite different cellular mechanisms^16^, landmark and dead reckoning pathways sum linearly on the compass network (Fig. 5b), with no evidence that one system is subservient to the other.

**Fig. 6.**
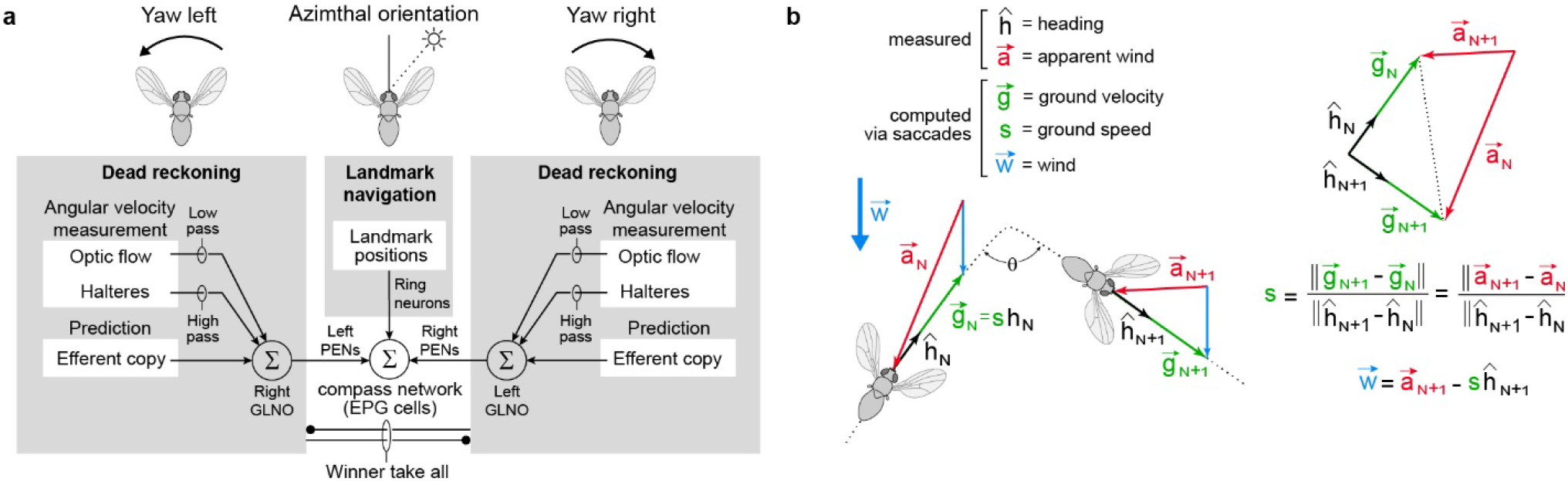
Feedback from the halteres during body saccades could be exploited to measure state parameters that would otherwise be unobservable. **a**, Schematic diagram a fly’s navigation system, summarizing major findings of this study. The landmark navigation system updates the internal compass by tracking the azimuthal position of external landmarks detected by the ring neurons. The dead-reckoning system incorporates angular velocity measurements from the visual system and halteres along with motor predictions to update the compass even when landmark cues are absent. **b**, Model of how flies could use their halteres to estimate ground speed 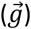 and wind speed 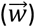, simplified from a prior analysis^48^ under the assumption that flies maintain constant ground speed before and after they saccade^43,62^. Flies cannot directly measure the absolute ground speed vector, but they can measure the apparent wind vector 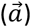 using their antennae^63,64^, and they know their heading vector 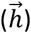 from the compass system. Ground speed, s, may be determined from the difference in the apparent wind vectors measured before and after a saccade, 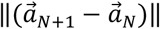, divided by the change in the heading 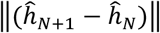. The wind vector 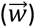 could then be calculated from the difference of the apparent wind and ground speed vectors. Using an iterative strategy, a fly could update its estimate of 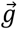 and 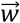 with each saccade.

In 1914, 21-year-old Lawrence Sperry won first prize at the Airplane Safety Competition in Bezon, France by demonstrating a gyroscopic stabilizer, the first practical autopilot^61^. With characteristic flair, Sperry flew his Curtis C-2 over the judges’ stand with hands raised off the flight controls and his co-pilot standing on one wing. In 1929, James Doolittle tested another of Sperry family invention—the gyrocompass—by making a 15-minute ‘blind flight’, relying solely on the instruments in his cockpit. We suspect that ancestral dipterans followed this same temporal sequence 250 million years ago. Halteres evolved first for stabilization reflexes, and then were subsequently incorporated into the compass system. Haltere-mediated reflexes allow flies to execute the sophisticated flight maneuvers such as landing upside-down and avoiding flyswatters, whereas haltere input to the compass endows them with greater navigational precision for dead-reckoning.

But why would a fly need to update its internal compass during a fast turn? Given that the mapping of the ring neurons onto the EPG population is stable from minutes to hours, a fly that deviates from its course due to perturbations could return to its original heading using landmark cues. Thus, the sensitivity to elevated angular velocities conferred by the halteres should not have a profound impact on directed navigation or menotaxis. Indeed, insects lacking halteres such as bees, moths, butterflies, and locusts are exceptional long-distance migrators^65^. Recent work, however, suggests haltere input to the CX might confer another advantage to flies that is linked to their characteristic habit of executing body saccades. Based on Kalman’s theory of observability, it has recently been proposed that the pattern of straight trajectories interspersed with rapid saccades constitutes a form of active sensing^46–49,66^. According to this hypothesis, sensory measurements made along two or more sequential straight flight segments provide a means of observing important state variables such as ground velocity, wind speed, and altitude—all of which would be unobservable by an animal flying on a straight trajectory at constant speed. A simple example of how an animal could render unobservable parameters observable by turning quickly is illustrated in Fig 6b. If a fly measures its heading and the apparent wind vector before and after a single saccade, it could calculate both its ground velocity and the wind speed provided it knows: (1) how far it has turned, and (2) possesses the circuitry for performing the requisite calculations. Evidence for these two criteria have now been met; whereas our results demonstrate how halteres provide a an estimate of heading changes during rapid turns, prior work shows that circuits within the Fan-Shaped Body can carry out vector computations^16,67,68^. The ability to make the two measurements in rapid succession—before and after a saccade—is advantageous because the relevant parameter such as wing speed will likely not have changed over the short interval. Further, the fly benefits from making its measurements while flying straight, as this reduces variability. The ability to accurately estimate ground speed, altitude, wind direction, and distance to objects would be particularly advantageous for odor localization tasks^69^. We propose that the halteres’ ability to measure fast angular velocities was a critical preadaptation for performing the computations necessary to estimate important state parameters with greater accuracy than other insects, thus leading to the evolution of saccades as a strategy for active sensing. If this hypothesis is correct, the advantages of this movement-based computation strategy conferred by the gyrocompass system may help explain unusual ecological diversity of the order Diptera.

## Methods

### Experimental subjects

All experiments were conducted on 3-to 5-day-old female *Drosophila melanogaster*, reared on standard cornmeal food at 25°C with a 12:12 hour (light:dark) cycle. For behavioral experiments, we used wild-type flies, descendants of a Heisenberg Canton-S stock (HCS). For functional imaging experiments, we generated flies expressing GCaMP7f^70^ in GLNO neurons by crossing the GAL4 driver line, w[1118]; P{y[+t7.7] w[+mC]=GMR76E11-GAL4}attP2) (BDSC_39933)^71^, to a GCaMP7f responder line, w+;UAS-tdTomato;UAS-GCaMP7f, which was constructed in our lab using w[1118]; P{20XUASIVS-jGCaMP7f}su(Hw)attP5; (BDSC_80906) and;; P{w[+mC]=UAS-tdTom.S}3 (gift from D. Anderson). To image EPG neurons, we crossed the split-GAL4 line, SS00096; (BDSC_86861)^59^, with the GCaMP7f responder line described above.

### Connectome path analysis

To find putative mechanosensory feedback pathways to the CX, we performed a polysynaptic connectivity search between wing or haltere campaniform sensilla afferents and GLNO neurons using FlyWire FAFB Materialization 783 (https://flywire.ai)^17,18^. We first identified neuron types that account for the top 75% of the total synapse count onto GLNO cells. We then identified the top 25% of neuron types that provide inputs to the members of this upstream population, thus identifying a set of possible bi-synaptic pathways to GLNO neurons. We then selected all neuron types receiving input from haltere campaniform afferents and forming output connections (above a synapse count threshold of 25) onto any member of the bi-synaptic input pathways to the GLNO cells described above (including the GLNO neurons themselves). For comparison, we performed an identical analysis using the wing campaniform afferents.

### Neurotransmitter identification

We deployed a genetic labeling strategy to identify the principal neurotransmitter released by GLNO neurons. We first constructed a split-GAL4 driver line, [76E11.AD; VT026017.DBD], that targets GLNO cells and crossed it to one of three reporter lines that we constructed to detect acetylcholine, GABA, and glutamate. Each line contains: (1) T2A-LexA::QFAD Trojan exon inserted in a MiMIC transposon site within the endogenous locus of a neurotransmitter gene (ChAT, GAD1, or vGLUT)^72^, (2) nuclear-targeted tdTomato under LexAop control to visualize brain wide neurotransmitter expression, and (3) GCAMP8m under UAS control to visualize the GLNOs. Brains of 3-to-5-day-old female adults were dissected at room temperature (∼20°C) in 1X phosphate buffered saline (PBS). Dissected brains were fixed in 4% paraformaldehyde diluted in 1X PBS for 25 min and washed 3 times for 15 min in 1X PBS. Brains were washed 5 times for 15 min in 0.3% PBS-T (1X PBS with 0.3% Triton-X), before being transferred to blocking solution (7.5% normal goat serum diluted in 0.3% PBS-T) for at least 1 hour at 20°C. The samples were then incubated with primary antibodies diluted in blocking solution (rabbit polyclonal anti-DsRED at 1:2000 and chicken anti-GFP 1:1000) for 4 hours at 20°C and 2 days at 4°C. Samples were subsequently washed 5 times for 15 min in 0.3% PBS-T and incubated with secondary antibodies (goat anti-rabbit Cy3 at 1:1500 and goat anti-chicken Alexa Fluor 488 at 1:1000, diluted in blocking solution) for 4 hours at 20°C and 2 days at 4°C. We mounted the samples on microscope slides with spacers using Vectashield Plus (VectorLabs) and imaged them on a confocal microscope (Zeiss LSM 880) with a 40X water immersion objective (1.6 zoom factor, 1-μm optical sections at 1-μm intervals, 1024×1024 pixel resolution).

### Simulating Coriolis forces

Nalbach’s micro-oscillation approach emerges from the basic physics of haltere motion. The sum of all forces acting on the end knob of the haltere in the body frame of the fly, ***F***_*h*_, is given by^3^:

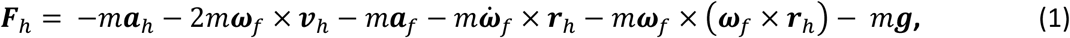

where *m*, ***r***_*h*_, ***v***_*h*_, ***a***_*h*_ are the mass, position, velocity, and linear acceleration of the haltere end knob; ***ω***_*f*_, 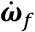 are the angular velocity and angular acceleration of the fly; ***a***_*f*_ is the linear acceleration of the fly; and **g** is the gravitational acceleration. The six terms on the right side of **Eq. 1** represent (in order): (1) the primary acceleration force (due to the oscillation of the haltere), (2) the Coriolis force (due to the linear velocity of the haltere on the rotating body of the fly), (3) the linear acceleration force (due to the linear acceleration of the fly), (4) the angular acceleration force (due to angular acceleration of the fly), (5) the centrifugal force, and (6) the gravitational force. These terms can be accurately calculated given the mass of the haltere, its kinematics in the fly frame, and the kinematics of the fly in the world frame. In a free-flying fly executing typical maneuvers, the four right-most terms are extremely small or can be safely ignored^1,3^, leaving only the so-called ‘primary force’ and the Coriolis force^1^, both of which vary in time as the halteres oscillate. The primary force results from the rapid oscillation of the haltere in its stroke plane and is present even when the fly is stationary, as in hovering flight or when the animal is rigidly tethered. These forces are relatively large and reside within the plane of haltere oscillation. The haltere is subjected to the Coriolis effect only when the fly’s body rotates while the halteres oscillate. The Coriolis forces are ∼10^-2^ times smaller than the primary forces, but have components that act perpendicular to the plane of oscillation, which is how the fly is thought to discriminate between the two^1^.

Given the basic physics of motion characterized by Eq. 1, the principle behind Nalbach’s approach is conceptually straightforward^3^. The third term in Eq. 1 represents the forces on the haltere resulting from the linear accelerations of the body, which are quite small during typical free flight behaviors. However, if the fly is tethered to a linear actuator (Fig. 2A), it is possible to impose oscillatory acceleration that replicates the time history of Coriolis forces that the animal would experience if it were rotating in free flight (Fig. 2C). A full derivation is provided in Supplementary Information, however a reasonable approximation for the acceleration amplitude, *A*, required to simulate an angular velocity, *ω*_z_, about the yaw axis is:

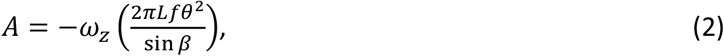

where *L* is the length of the haltere (∼290 μm), *f* is the stroke frequency of the haltere (which is the same as the wingbeat frequency, ∼200 Hz), *θ* is the amplitude of haltere oscillation (∼80°), and *β* is the angle at which the haltere stroke plane is inclined caudally with respect to the bilateral body axis of the fly (∼30°). The fly is oscillated using a sine wave of amplitude, A, and frequency, 2f, a doubling that is necessary to force the haltere into a ‘figure-of-eight’ in each stroke, thus simulating the Coriolis forces experienced during yaw^1,3^. Using the values listed above, Eq. 2 indicates than an angular velocity of 1000 ° s^-1^ may be simulated by a harmonic oscillation with an amplitude of 4 μm and a frequency of 2f. By choosing the direction, frequency, and phase of linear oscillation, it is also possible to accurately simulate roll and pitch, as well as any arbitrary rotation in the yaw-roll plane. See Supplementary Information for more details.

The Flybratron consists of a voice coil actuator controlled by an ItsyBitsy microprocessor (Asafruit), with a 5G MEMS accelerometer for measuring the oscillations of the fly. We used a custom-built optoelectronic wingbeat analyzer^19^ to track wing motion in real-time and generate a phase pulse adjusted to occur precisely at the start of the haltere stroke^19,73^. Each wingbeat, the microprocessor generates a harmonic function with an amplitude and frequency that simulates the targeted value of angular velocity. The actuator for oscillating the fly consists of an 8 ohm, 20W voice coil oscillator (Hilitand) driven by a 20W digital audio amplifier (Kinter, K3118). A 50 mm long, 6 mm diameter polished stainless steel rod was firmly bolted to the face of the voice coil, supported by a 20 mm long linear bearing that restricted its motion to one dimension, perpendicular to the face of the voice coil. The voice coil and bearing were bolted to a custom-built, 3D-printed plastic housing designed for that purpose. Another small 3D-printed part fixed to the end of the stainless-steel rod held a 3-axis, 5 G accelerometer (Analog Devices) and the female D-Sub connector into which each tethered fly was placed. The entire apparatus was attached to a 3-axis micromanipulator so that the position of the fly could be adjusted.

To control the Flybratron, we designed a printed circuit board that housed the microcontroller together with ancillary electronics and connectors. The inputs to the board were the synch pulse from the wingbeat analyzer, output signals from the accelerometer, and serial commands from the Python script running each experiment. The board provided an output signal to drive to the voice coil amplifier. The firmware installed on the device generates the pattern of motion necessary to simulate the Coriolis forces experienced by the fly during rotations about any axis at any specified angular velocity. Because the frequency of the phase-locked wings and halteres can change over time, the firmware was programmed to recalculate the required oscillation waveform on a stroke-by-stroke basis. As documented more fully in Supplementary Information, simulating roll and yaw requires oscillating the fly perpendicular to its long axis at a frequency that is once (for roll) or twice (for yaw) the haltere stroke frequency. Simulating pitch requires oscillating the fly parallel to its long axis at the haltere stroke frequency.

### Tethering flies for behavioral experiments

Wild-type flies were anaesthetized on a 4℃ cold plate and then attached to a custom-built holder at their notum using UV-curing glue (Bondic). The holder consisted of 220 μm tungsten rod (AM systems) glued inside a 10 mm long, 0.55 mm diameter stainless steel tube which was then soldered into a male D-Sub connector (DigiKey). The tungsten rod extended from the stainless-steel tube by 3 mm and the entire holder was 25 mm in length from the tip of the D-sub connector to the tip of the tungsten rod. Typically, we would tether 4-6 flies in one session and then provide them with tiny (∼2 x 2 mm) piece of paper to hold with their legs to inhibit flight until the start of the experiment. The flies were placed in the apparatus by inserting the holder into a female D-Sub pin fixed to the end of the Flybratron.

### Phase calibration

The wingbeat analyzer generates a stable synch pulse each stroke just before the end of the downstroke (Fig. 2b). The delay between this pulse and the start of the haltere downstroke could differ a small amount across preparations due to variation in the precise placement of each fly in the optical pathway between the LED and the sensor of the wingbeat analyzer. To correct for this variation, we performed a brief calibration procedure prior to data collection for each fly. First, the fly was aligned by eye within the optical axis of the wingbeat analyzer while inspecting both the voltage output representing the wing shadows from both wings as well as the live video image from Kinefly. After alignment, the magnitude of the left and right wing signals were balanced when the video image indicated that the fly was trying to fly straight, and adjusted to ensure the signals did not saturate when stroke amplitude was large. Next, we ran a calibration script in which the Flybratron runs a series of simulated yaw rotations at 1000 ° s^-1^ to the left and right while introducing a series of pre-specified phase advances and delays between the synch pulse from the wingbeat analyzer and the start of each oscillation cycle of the Flybratron. By inspecting the behavioral responses of the fly at different phase values, we selected the phase delay that evoked steering responses of equal and opposite magnitude evoked by simulated yaw rotations to the left and right. In all cases, the phase delay that evoked the best response was very close to what was expected given the expected temporal position of the wingbeat analyzer synch pulse. Further, as long as we aligned flies within the optical axis of the wingbeat analyzer carefully, the calibration procedure yielded an optimal phase-delay value that was nearly identical for each individual.

### Mounting flies to the flight stage for functional imaging

To stimulate the halteres during functional imaging, we mounted each fly by its head to a custom-built stage as previously described^74^, but with the thorax attached separately to the Flybratron by a fine, horizontal tungsten wire glued to the anterior notum (Fig. 3a,b). During the tethering process, we used two neodymium magnets (K&J Magnetics) to firmly, but temporarily, secure the tungsten wire to the flight stage. A 12.7 x 3.2 x 1.6 mm bottom magnet was permanently glued to the tungsten wire; a parallel to 6.4 x 3.2 x 1.6 mm top magnet was placed on the top surface of the flight stage to hold the bottom magnet and tungsten wire in place while the fly was tethered. The preparation was then moved to a custom-built laser cutter (https://github.com/willdickson/flasercutter), which scored a trapezoidal path on the back of the head capsule. After covering the back of the head with physiological saline, the trapezoidal section of cuticle was removed with fine forceps, providing imaging access to the appropriate brain region. After transferring the preparation to the 2-photon microscope, we bonded the bottom magnet (with the tungsten rod and fly attached) to the end of the Flybratron using UV-cure adhesive (Bondic), after which the retaining magnet on top could be safely removed. Prior to imaging, we carefully translated the Flybratron ∼15 µm posteriorly and ventrally to ensure that the tungsten wire was not in contact with the bottom of the flight stage during experiments.

The Flybratron firmware requires a stable, phase-locked pulse each and every wingbeat in order to create and deliver the oscillation waveform that simulates the Coriolis forces on the halteres. Because of insufficient clearance for a wingbeat analyzer, we designed and implemented a simple beam-break device, consisting of a 1mW, 780nm laser (VLM-780, Quarton Inc.) and a photodiode sensor (OPT101, Texas Instruments) aligned so that the wings would break the light path once each cycle when they were maximally elevated at the upstroke-to-downstroke reversal. Because the wings are almost perfectly in phase at this time in the stroke, the system registered a single beam-break event. The analog voltage signal from the photodiode was conditioned on-line using a custom-built 4-pole, low-pass Bessel filter with a cut-off frequency of 2kHz. We then fed this processed signal into a digital stimulator (PG4000, Neuro Data Instruments) to generated a brief TTL pulse suitable for the Flybratron firmware.

### 2-photon recording

We recorded GCaMP7f and tdTomato fluorescence in tethered, flying flies using a two-channel, 2-photon microscope (Thorlabs) equipped with a galvo-galvo scanner as previously described^31,74^. In most experiments, we used a 40x Nikon CFI160 Apo NIR, water-immersion objective (0.8 N.A., 3.5 mm W.D.). For GLNO recordings, we captured 40.5 x 27 µm images with 96 x 64 pixel resolution at 15.7 Hz. Because the Protocerebral Bridge is not restricted to a flat horizontal plane, we used a piezo-ceramic linear objective drive (P-726, Physik Instrumente GmbH and Co. KG) to image three x-y planes separated by 14 µm along the z axis. The time required to scan three image planes and the requisite fly-back time resulted in an effective scan rate of 9.5 Hz. For the EPG recordings in Fig. 4, which required a higher temporal resolution, we used 10x Nikon CFI160 Plan Fluor, water-immersion objective (0.3 N.A., 3.5 mm W.D.). The lower numerical aperture results in a thicker imaging plane, obviating the need to image multiple planes and allowing a scan rate of 38 Hz. To reduce light pollution, the microscope was housed in a light-tight enclosure and the visual arena panels (see below) were covered with transmission filters^74^. We tracked left and right wingstroke amplitude with Kinefly^75^ at 100 Hz, which introduced a 20 ms image processing delay that we corrected off-line. The wings were illuminated using four horizontal fiber-optic IR light sources (M850F2, Thorlabs) distributed in a 90° arc behind the fly.

At the start of each recording session, we selected a field of view for data collection centered along the bilateral axis based on tdTomato expression. The Regions of Interest (ROIs) that were used for quantifying fluorescence were determined during off-line data analysis. To correct brain motion in the horizontal plane, we registered both fluorescence channels for each frame using the cross-correlation between the tdTomato image and the trial-averaged image. To account for motion along the vertical axis, we normalized GCaMP7f fluorescence to tdTomato fluorescence. To standardize the time-varying neuronal signal for each individual fly, we normalized baseline-subtracted fluorescence in each ROI. For GLNO neurons, we normalized to the baseline magnitude: ΔF/F = (F_t_ – F_0_) / F_0_; For EPG neurons, we normalized to the maximum observed: ΔF/F = (F_t_ – F_0_) / (F_t_ – F_0_) _max_, where F_t_ is the time-varying neuronal signal and F_0_ is the baseline signal. F_0_ was defined as the 5^th^ percentile ROI pixel fluorescence in each trial, after subtraction of the fluorescence of the 10% dimmest pixels within the entire FOV for every frame.

To represent the azimuthal compass direction in the brain, we calculated the activity bump of EPG neurons in the Ellipsoid Body reference frame^50^. Bump position was calculated for every imaging frame from the circular mean of glomerular ΔF/F in EB coordinates, based on anatomical connectivity between the PB and EB^76^. The ROIs that were used for quantifying fluorescence were determined during off-line data analysis. For the GLNO neurons, the ROIs were determined by thresholding the terminal region of the cells within in the noduli (Fig. 3d). For the EPG cells, we manually assigned a reference region of interest (ROI_ref_) to each PB glomerulus with EPG neuron innervation. To avoid signal pollution from neurites or somas of neighboring EPG cells, only pixels within the glomerulus that covaried with the GCaMP7f fluorescence within ROI_ref_ were included in the ROI for each glomerulus.

### Presentation of sensory stimuli during functional imaging

Visual stimuli were presented on a 96 x 32 array of LEDs^77^ that covered 216° of azimuth and 74° of elevation. Each trial began with a closed-loop epoch in which the angular velocity of a pattern consisting of two bright 9°-wide vertical stripes was determined by the L-R wingstroke amplitude measured in real-time using Kinefly. We used two stripes (positioned 180^°^ apart) instead of one, because this promoted bouts of stripe fixation on a display with limited azimuthal width (216°). The haltere and visual stimuli simulating self-rotation were presented in open-loop at pre-determined angular velocities and durations. Open-loop haltere stimuli were presented via the Flybratron as described above. Open-loop visual stimuli consisted of one of three patterns: (1) a pattern of randomly spaced bright columns of variable width (resembling a barcode), (2) a single bright 9°-wide bright vertical stripe, or (3) a 2-by-2 pixel ersatz sun presented at 15° inclination relative to the fly’s anteroposterior head axis. Each open-loop rotational stimulus was flanked by 1 s periods in which a control pattern was displayed. For experiments in which we presented a rotating barcode pattern, the control pattern consisted of a static barcode. For experiments in which we presented a rotating landmark (bright spot or stripe) or haltere stimulation alone, the control pattern was a dark, blank display. Open-loop presentations were interspersed with 4 s epochs of closed-loop stripe fixation. To reduce luminance changes, the closed-loop stripe was dark-on-bright during barcode trials and bright-on-dark during stripe or sun trials. Every open loop condition was presented to the left and to the right with pseudo-randomized order.

### Saccade detection

To detect saccades during 2-photon imaging, we filtered the voltage output of Kinefly encoding the bilateral difference in wingstroke amplitude (sampled at 100 Hz) and conditioned it with a real-time, first-order, 2-6 Hz band-pass filter. We then set a threshold of 12^°^ for the filtered L-R WSA signal. For trials in which we triggered a simulated body rotation when the fly executed a saccade, we measured a processing latency of 20 ms between the peak of the saccade and the start of the haltere stimulus. Our software enforced a refractory period of 1 s after each saccade detection event (and waited until the L-R WSA signal fell below 7.2°) to ensure that we did not subject the flies to rapid bursts of successive haltere stimulation.

### Statistics and Reproducibility

All experiments were analyzed with custom software written in Python. Sample sizes refer to the number of individuals tested and are provided for every experiment in the figure panels. For all experiments we calculated a single mean time-history trace for each individual fly by averaging the data from individual trials after aligning the data to stimulus onset. We then constructed grand means across flies. To quantify variance across individuals, we determined the boot-strapped, 95% confidence intervals of the individual means (CI). The probability that any particular data value or trace would fall with the distribution of another set of measurements may be inferred directly from the inspection of the 95% CI domain.

## Data and materials availability Data availability

The data and supporting the findings of this study will be made available in a public repository upon publication.

## Acknowledgements

We wish to thank Kevin Mills, Haoru Li, and Anne Erickson for their comments on this manuscript. The research reported in this publication was supported by the National Institute of Neurological Disorders and Stroke of the NIH (R01NS136988).

## Author Contributions

Will Dickson designed all the hardware and software components of the Flybrator and wrote the Supplementary Information. Ruoxi Wang collected and analyzed all the data presented in Fig. 2. Ivo Ros collected and analyzed all the data presented in Figs. 3-5 and performed the connectomic analysis in Fig. 1d. Jaison Omoto conducted all the required genetic crosses and performed the transmitter identification of the GLNO neurons. All authors collaborated on the figures. Michael Dickinson conceived of the project, collaborated on planning the experiments, and wrote the first draft of the manuscript.

## Competing of Interest

The authors have competing interests to report.

## Extended Data Figures

**Extended Data Figure1.**
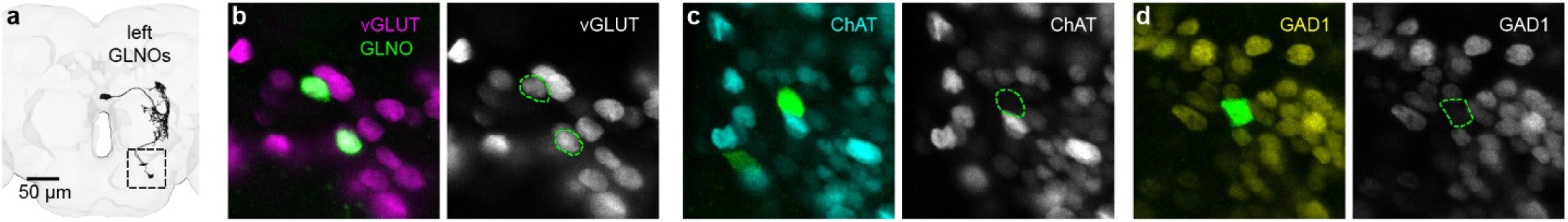
Evidence that GLNO neurons are glutamatergic. **a**, Anterior view of left GLNO neuron morphologies in FAFB-FlyWire. Dashed box indicates approximate region in b-d. **b**, Left: Merged confocal images with RFP (green) in GLNO neurons under Gal4 control and GFP (magenta) expressed using a Trojan-LexA for the glutamate marker vGlut. Right: vGlut labeling (grey) with the soma of a the GLNO cells outlined in dotted green lines. **c**, As in b, but GFP indicates an acetylcholine-specific LexA reporter associated with the ChAT gene (cyan, left; grey, right). **d**, As in b, but GFP indicates a GABA-specific LexA reporter associated with the gad1 gene (yellow, left; grey, right).

## Supplementary Video Legends

**Supplementary Video 1. Behavioral responses to simulated yaw rotation using the Flybratron**.

A tethered fly is subjected to a series of simulated yaw rotations via micro-oscillation, comparable to the data presented in Fig. 2e. The beating halteres are clearly visible just behind the base of the wings and in front of the mesothoracic legs. Bottom panel: Wing motion (L-R WSA, dark trace) in response to 0.25 s rotational stimuli (grey trace), presented in both directions and at increasing angular velocities. Fly thorax oscillations (bottom trace) are too small to be seen in the top video sequence. In addition to the large changes in wing stroke amplitude, the neck, leg, and abdomen motor systems also visibly respond to the yaw stimuli in a directional manner. Note the sustained response in wing motion following presentation of the first stimulus in each stimulus pair, which is evidence for the presence of integral feedback in the haltere-motor system. Playback speed is 1x real time.

**Supplementary Video 2. 2-photon imaging of GLNO neurons during simulated yaw rotation**.

A flip-flop pattern of GLNO activity elicited by micro-oscillation simulating alternating left and right yaw rotations. Both GLNOi and GLNOii neurons express GCaMP7f. See Fig. 4a for recording arrangement. Simulated yaw rotations to the left (top trace) activate the left GLNOs; yaw right activates right GLNOs. The bottom trace represents compensatory wing motion responses (L-R WSA). Note that Flybratron oscillations do not cause any visible movement artifact during stimulus presentation. Playback speed is 1x real time.

**Supplementary Video 3. 2-photon imaging of the EPG neurons in the Protocerebral Bridge during simulated yaw rotation**. As the fly is subjected to yaw rotations to the left and right, the coordinated bumps of EPG activity in the left and right halves of the Protocerebral Bridge move in the appropriate direction, opposite that of the simulated rotations. The magnitude of EPG bump movements increases with the angular velocity of the yaw rotation. The trace below indicates the time course and angular velocity of the stimuli. Playback speed is 1x real time.

## Supplementary Information

### Simulating Haltere Coriolis Forces Using Linear Oscillations

#### Forces on the haltere

Nalbach and Hengstenberg [1] derived a complete expression for the total inertial and gravitational forces acting on the haltere’s end knob in body frame coordinates:

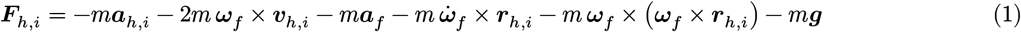

where *m*, ***ω***_*f*_, ***v***_*h,i*_ and ***a***_*h,i*_ are the mass, angular velocity, velocity and linear acceleration of the haltere end knob; ***a***_*f*_ is the linear acceleration of the fly; and *i* indicates the anatomical side of the haltere attachment, (R = right, L = left).

The six terms on the right side of Eq. 1 represent (in order): (1) the primary acceleration force (due to the oscillation of the haltere), (2) the Coriolis force (due to the linear velocity of the haltere on the rotating body of the fly), (3) the linear acceleration force (due to the linear acceleration of the fly), (4) the angular acceleration force (due to angular acceleration of the fly), (5) the centrifugal force, and (6) the gravitational force. Under most free flight conditions, the four right-most terms are extremely small or can be safely ignored[2] leaving only the primary acceleration force and the Coriolis force, both of which vary in time as the halteres oscillate. However, only the Coriolis term is sensitive to the angular velocity of the fly. Nalbach’s approach for simulating rotation is to subject the fly’s body to a linear acceleration that precisely mimics the time history of the Coriolis forces acting on the halteres during a free flight rotation. Thus, the relevant equation reduces to:

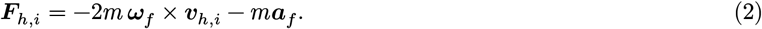

The first term in Equation 2 represents the Coriolis force acting on the haltere’s end knob:

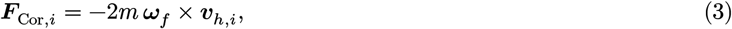

and second term in is the effect of the fly’s linear body acceleration on the end knob of the haltere,

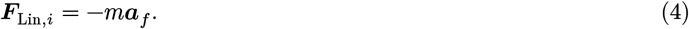

#### Relating angular velocity to linear acceleration

In this approach, a fly is rigidly fixed to an actuated tether connected to a voice coil oscillator that provides linear acceleration along the *x, y*, and *z* axes while remaining rigid about all rotational axes. In the following description, the orientation of the *x, y* and *z* axes relative to fly’s body are assumed to be as shown in Figure 1. The 3D fly body model used is this figure was adapted from the model made available by Taraga et al. [3]

**Figure 1:**
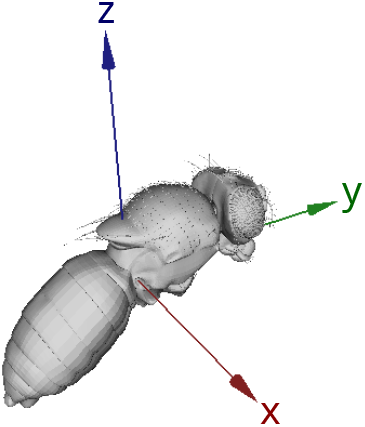
Illustration showing the location of the *x, y*, and *z* axes relative the body of the fly.

As the fly is rigidly tethered and does not rotate relative to the laboratory frame, the actual angular velocity, and consequently the true Coriolis force in Equation 2, is zero. Under these conditions, the only non-zero term is the force resulting from linear body acceleration. The linear acceleration required along the *x, y* and *z* axes to simulate a specific angular velocity vector ***ω***_*f*_ must satisfy the following:

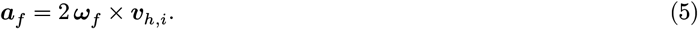

Expanding the vectors in Equation 5 into their respective components and expressing the cross product as a matrix-vector multiplication yields:

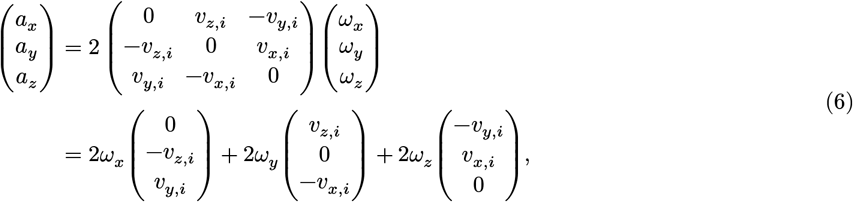

where

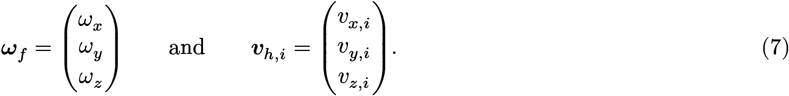

Equation 6 demonstrates how the linear acceleration components necessary to simulate a given angular velocity depend on both the components of that angular velocity and the instantaneous haltere velocity.

Following Nalbach and Hengstenberg [1], the acceleration acting on the haltere can be decomposed into three orthogonal components. A radial component within the haltere stroke plane pointing outward, a tangential component within the haltere stroke plane aligned with the instantaneous haltere velocity, and a perpendicular component normal to the haltere stroke plane directed rostrally.

Based on the expected directional sensitivities of the campaniform sensilla fields at the base of the haltere [1,2], only the component of the Coriolis force perpendicular to the haltere stroke plane is detected by the fly. With this constraint, the expression in Equation 5 may be simplified by projecting it onto the rotational axis, ***h***_*i*_, of the haltere:

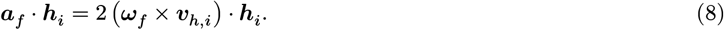

Equation 6 then reduces to the following scalar form:

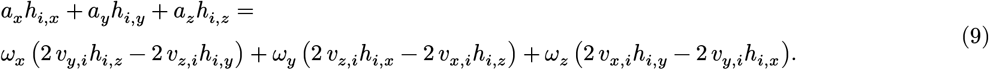

Note, that the components of angular velocity, ***ω***_*f*_, and linear acceleration, ***a***_*f*_, are common to the entire body, whereas the haltere end knob velocity, ***v***_*h,i*_, and rotation axis, ***h***_*i*_, are specific to the left or right side, *i*, of the fly. Thus, to simulate the Coriolis forces for arbitrary angular velocities, expressions for the left and right end-knob position, velocity and haltere rotation axes are derived.

#### Haltere kinematics

In this section expressions for haltere kinematics that incorporate the key anatomical features are derived. Again following the work of Nalbach and Hengstenberg[1], it is assumed that the haltere’s motion is restricted to a vertical stroke plane tilted caudad by an angle *β* ≈ 30° (Figure 2). The instantaneous angular position of the haltere, *θ*(*t*), within this plane (Figure 3) is approximated as a sinusoid of the form:

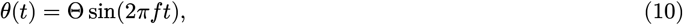

where *f* is frequency and Θ is the amplitude.

The position of the end knobs of the right and left halteres, ***r***_*h*, R_ and ***r***_*h*, L_, can then be specified as follows:

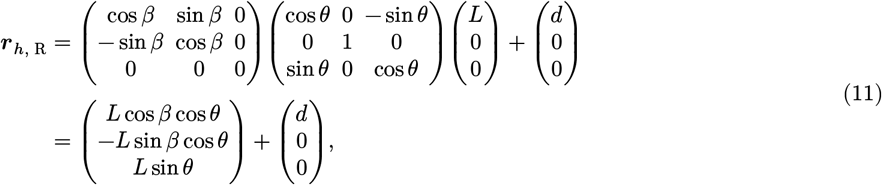

and

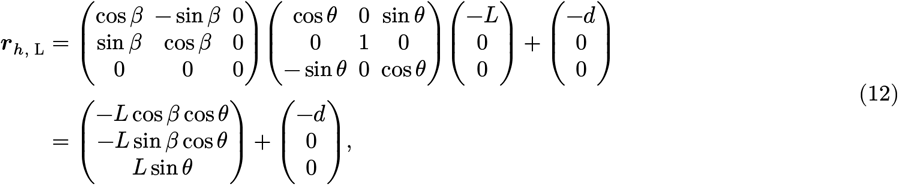

where *β, L, d* are the tilt angle of the haltere (cuadad), the length of the haltere (from base to end knob), and the distance from the haltere base to the fly’s center of mass.

Taking the derivative of ***r***_*h*, R_ and ***r***_*h*, L_ with respect to time gives the velocity vectors, ***v***_*h*, R_ and ***v***_*h*, L_, for the haltere end knobs:

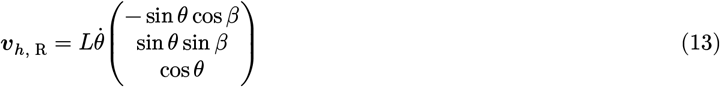

and

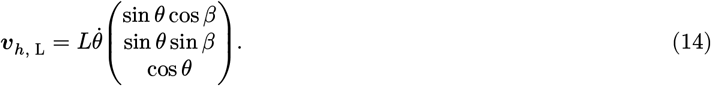

Comparing the vector components of Equations 13 and 14 shows that the *x* components of the right and left haltere velocities have the same magnitude and opposite sign, whereas the *y* and *z* components have the same magnitude and the same sign:

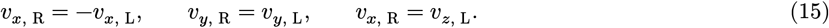

Expressions for the rotation axes of the right and left halteres, ***h***_R_ and ***h***_L_, may be derived by starting with a unit vector pointing down the *y*-axis and rotating it caudad by the angle *β*, to either the right or left:

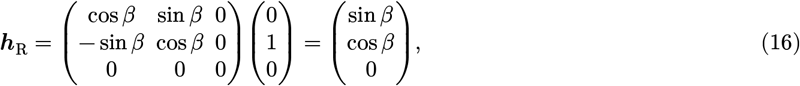

and

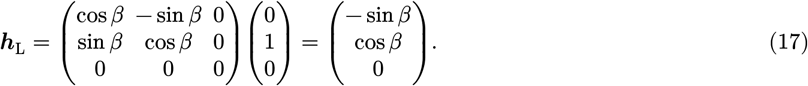

**Figure 2:**
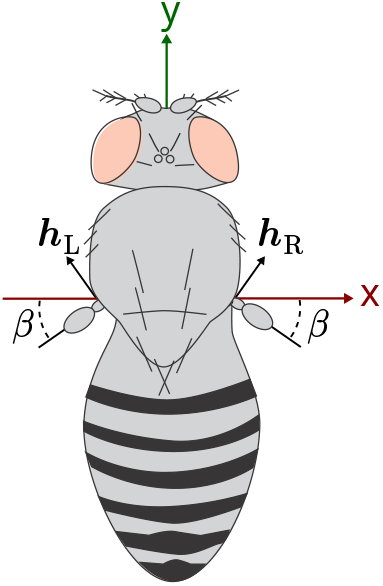
Top view of fly illustrating the tilt angle, *β*, of the haltere stroke plane relative to the body of the fly and the haltere rotation axes ***h***_R_ and ***h***_L_.

**Figure 3:**
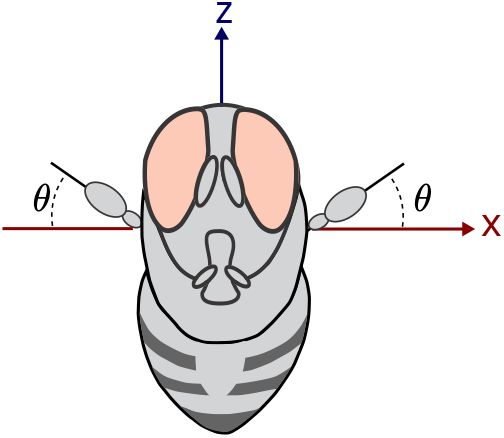
Front view of fly illustrating stroke position angle *θ*.

#### Simulating angular velocity

In this section, expressions relating the components of the angular velocity to the components of the linear acceleration required to simulate body rotation when the fly is attached to a rotationally rigid tether are developed for the right and left halteres.

The expression relating the components of linear acceleration to angular velocity vector, Equation 9, may be simplified using the haltere axes (Equations 16 and 17). Because the *z* component of the haltere axes are always zero, Equation 9 reduces to:

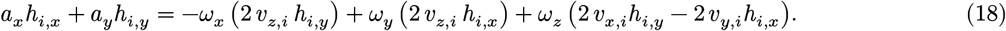

Substituting the components of the right haltere axis, Equation 16, into Equation 18 generates an expression relating linear acceleration to angular velocity for the right haltere:

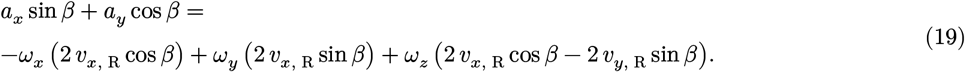

Similarly, substituting the components of the left haltere axis, Equation 17, into Equation 18 yields an expression for the left haltere:

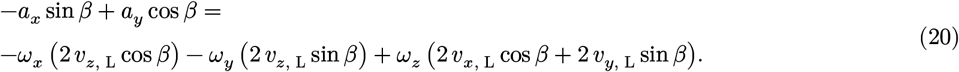

Next, using the fact, established in Equation 15, that the *x* components of the left/right haltere velocities have the same magnitudes and opposite signs whereas the *y* and *z* components have the same magnitude and sign, Equation 19 for the right haltere, may be rewritten as:

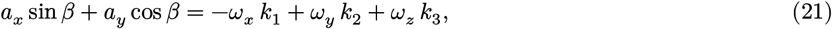

In a similar manner, Equation 20 for the left haltere, may be written as:

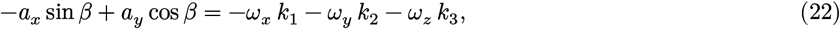

where

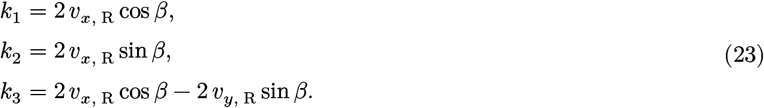

The simulation of pitch, *ω*_*x*_, requires that the left and right halteres undergo accelerations of the same sign. This symmetric acceleration requires linear acceleration, *a*_*y*_, along the *y*-axis. Conversely, roll and yaw (*ω*_*y*_ and *ω*_*z*_) require accelerations of opposite signs between the two sides, which may be generated by linear acceleration, *a*_*x*_, along the *x*-axis.

Furthermore, in the scalar expressions for the left and right Coriolis accelerations (Equations 21 and 22), the coefficient *k*_2_ multiplying the roll component, *ω*_*y*_, is proportional to sin *β*. Consequently, the Coriolis acceleration for roll is only non-zero if the tilt angle, *β*, is non-zero. In addition, since the *x*-component of the linear acceleration also appears as a multiple of sin *β*, it follows that neither roll nor yaw can be effectively simulated without a non-zero tilt of the haltere stroke plane.

#### Yaw case

For a pure yaw rotation, *ω*_*x*_ = 0 and *ω*_*y*_ = 0. Because the accelerations for the two sides must be of opposite signs, the simulation can only be achieved using linear acceleration along the *x*-axis, *a*_*x*_, thereby reducing the system to a single equation:

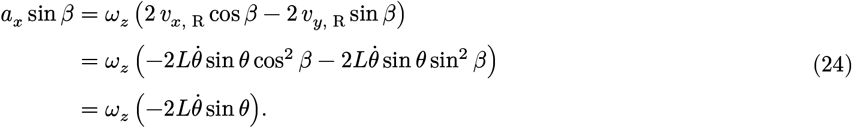

Solving for *a*_*x*_ yields the linear acceleration required to simulate the angular velocity *ω*_*z*_:

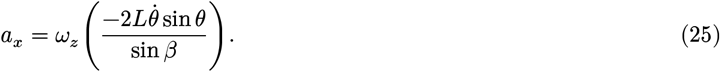

Thus, simulating yaw requires application of a time-varying linear acceleration along the *x*-axis that is proportional to 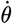. Here, *L* and *β* are morphological parameters whereas *θ* and 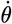, which represent the position and velocity of the haltere, are periodic functions that depend on the flapping frequency, *f*, and stroke amplitude, Θ, of the haltere.

If *θ* is a sine wave, as given in Equation 10, then 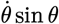 may be rewritten as a Fourier series:

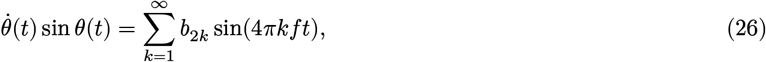

where

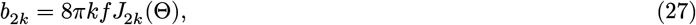

where 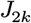 is the Bessel function of the first kid of order 2*k* (see Supplementary appendix for a complete derivation). Using this result, the expression for the linear acceleration required to simulate yaw in Equation 25 may be rewritten in terms of its frequency components as:

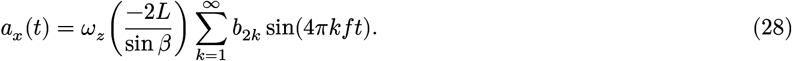

Truncating the Fourier series at the first term yields a useful approximation for *a*_*x*_(*t*):

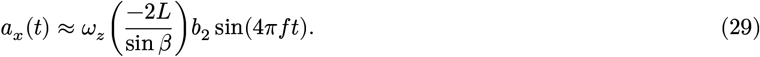

This approximation consists of a single sine wave at twice the haltere flapping frequency:

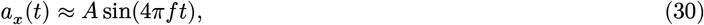

with an amplitude given by:

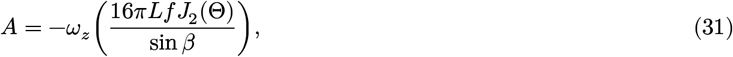

For small Θ, the Bessel function *J*_2_ may be approximated as:

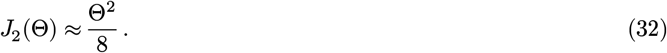

With this approximation the amplitude of the first Fourier component of the required linear acceleration becomes:

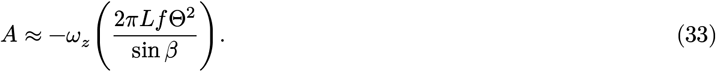

The difference between these two approximations (Equation 31 and Equation 33) may be illustrated by considering the ratio the amplitudes *A*_Fourier_ and *A*_Small Θ_:

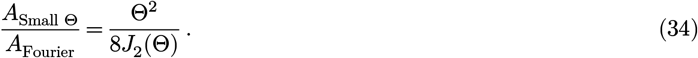

In designing and implementing our experiments, the acceleration of the haltere was approximated using a sign wave, as given by Equation 30, and the simplified expression for given by Equation 33 was used for calculating the amplitude of the oscillations.

#### Roll case

For a pure roll rotation, *ω*_*x*_ = 0 and *ω*_*z*_ = 0. As with yaw, the accelerations on both sides of the fly must be of opposite; therefore, the simulation is achieved using linear acceleration along the *x*-axis:

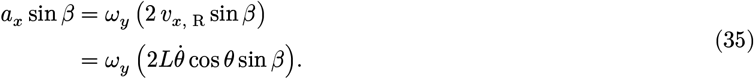

Solving for *a*_*x*_ yields:

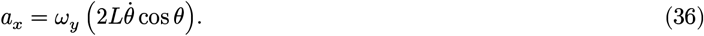

Thus, simulating roll requires a time-varying linear acceleration along the *x*-axis proportional to 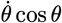. If *θ* is a sine wave, as given in Equation 10, then 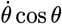 can be rewritten as Fourier series:

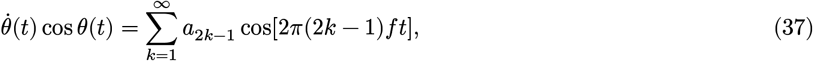

where

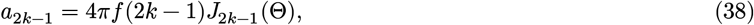

and 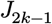 is the Bessel function of the first kind of order 2*k* − 1 (see Supplementary appendix for a complete derivation). Using this result, the expression for the linear acceleration required to simulate roll in Equation 36 may re-written in terms of its frequency components as:

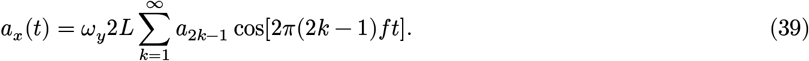

As with yaw, truncating the Fourier series at the first Fourier component yields a useful approximation for the linear acceleration, *a*_*x*_(*t*):

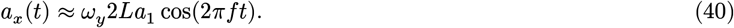

This approximation then consists of a single cosine at the haltere flapping frequency:

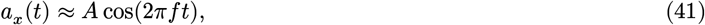

with amplitude:

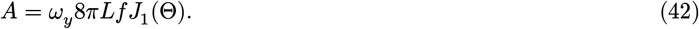

For small Θ, the Bessel function *J*_1_ may be approximated to leading order in Θ by:

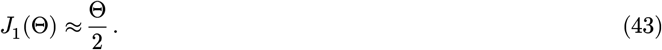

With this approximation, the amplitude of the first Fourier component of the required linear acceleration becomes:

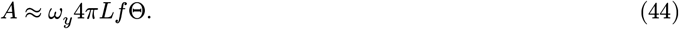

The difference between these two approximations can be illustrated by considering the ratio the amplitudes the sinewaves, *A*_Fourier_ and *A*_Small Θ_, given in Equation 42 and Equation 44 as follows:

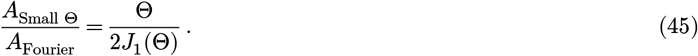

In designing and implementing our experiments, the acceleration of the haltere was approximated using a sign wave, as given by Equation 41, and the simplified expression for given by Equation 44 was used for calculating the amplitude oscillations.

#### Pitch case

For a pure pitch rotation, *ω*_*y*_ = 0 and *ω*_*z*_ = 0. In this case, the accelerations for the two sides must have the same sign. The simulation is therefore achieved using linear acceleration along the *y*-axis, *a*_*y*_ such that:

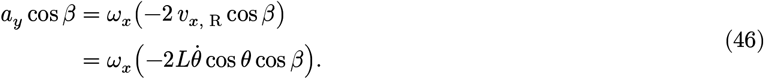

Solving for *a*_*y*_, the required linear acceleration is:

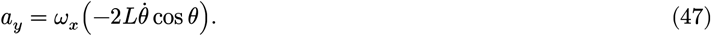

Therefore, to simulate pitch, a time-varying linear acceleration along the *y*-axis proportional to 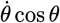 must be applied. Similar to the expression for roll, Equations 37 and 38 can be used to rewrite the required linear acceleration in terms of its frequency components:

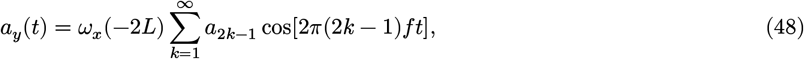

where *a*_2*k*−1_ is given by Equation 38.

As with yaw and roll, truncating the Fourier series at the first Fourier component yields a useful approximation for the linear acceleration, *a*_*y*_(*t*):

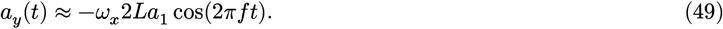

This approximation then consists of a single cosine at the haltere flapping frequency

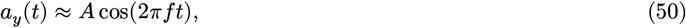

with amplitude given by:

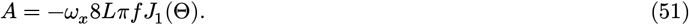

Similar to yaw and roll, using the approximation for the Bessel function *J*_1_ in Equation 43 for small Θ, first Fourier component of the required linear acceleration becomes:

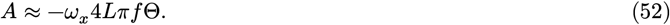

The difference between these two approximations (i.e. Equation 51 and Equation 52) may be illustrated by considering the ratio the amplitudes the sine waves, *A*_Fourier_ and *A*_Small Θ_:

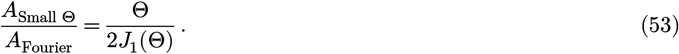

In designing and implementing our experiments, the acceleration of the haltere was approximated using a sign wave, as given by Equation 50, and the simplified expression for given by Equation 52 was used for calculating the amplitude oscillations.

#### Oscillation amplitude

Equations 33, 44 and 52 provide expressions for the acceleration amplitudes required to simulate yaw, pitch, and roll angular velocities. Integrating these accelerations, Equations 30, 41 and 50, twice yields the corresponding displacement amplitudes. Figure Figure 4 plots these amplitudes for angular velocities from 0 to 2000 (°/*s*). The plot assumes typical anatomical properties for the halteres: the length *L* = 290*μ*m, flapping frequency *f* = 200 Hz, and haltere stroke amplitude Θ = 90°.

**Figure 4:**
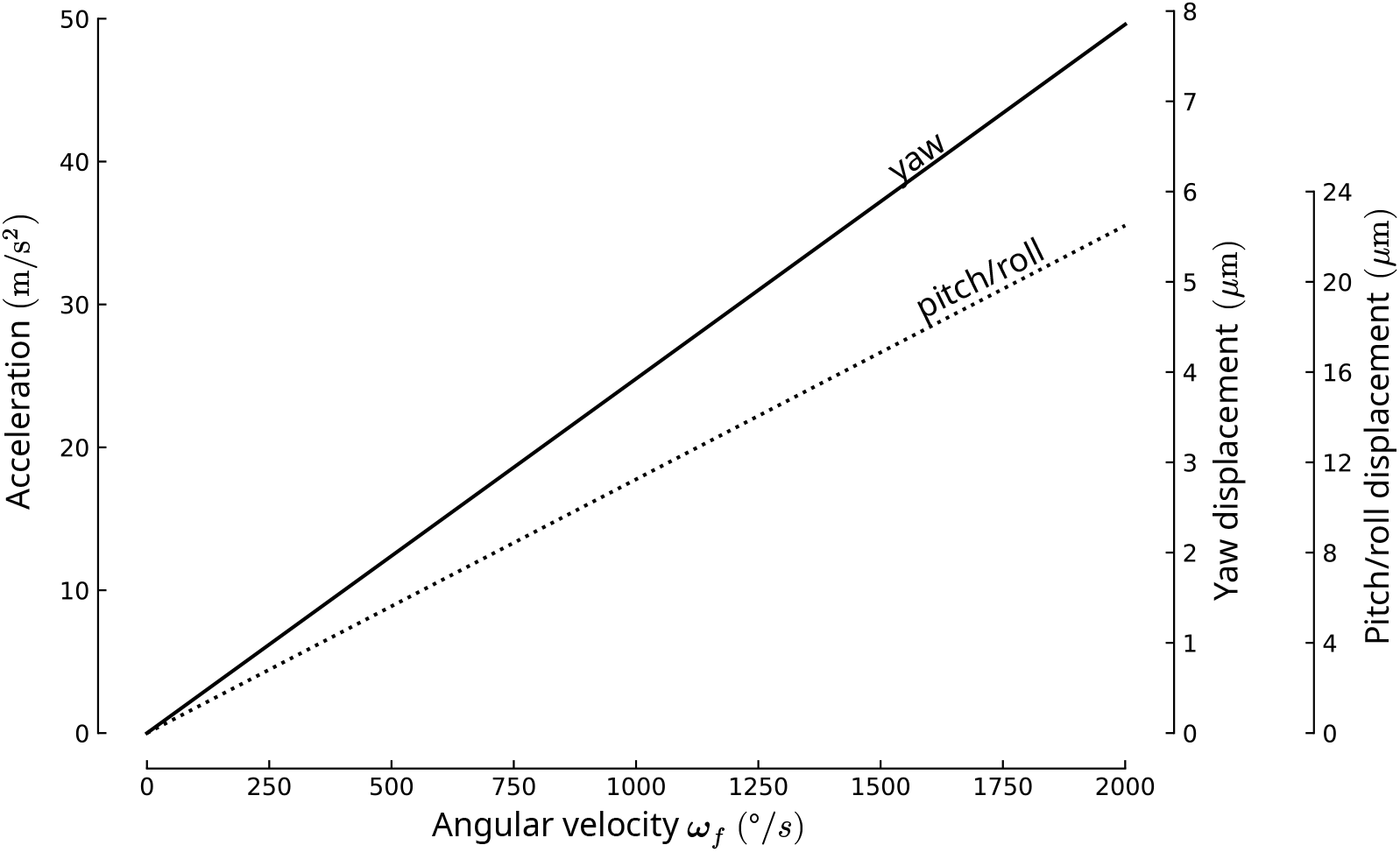
Acceleration and displacement amplitudes as a function of yaw, pitch and roll angular velocity.

## Supplementary appendix

In this appendix the Fourier series for the two functional forms of linear acceleration required to simulate roll, pitch, and yaw are derived:

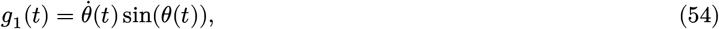

and

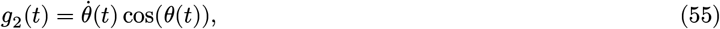

where the stroke angle, *θ*(*t*), is a sinusoid with frequency, *f*, and amplitude, Θ, as defined in Equation 10. A periodic function *g*(*t*) with frequency *f* may be expanded into a Fourier series as follows:

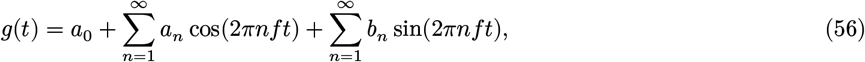

where the cosine and sine coefficients *a*_*n*_ and *b*_*n*_ are given by:

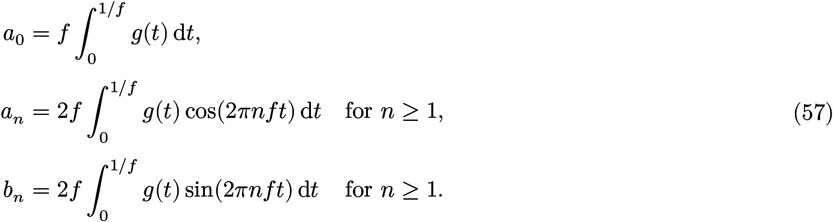

By applying chain rule and using the fact that *θ*(*t*) = Θ sin(2*πft*) the expression for *g*_1_(*t*) can be rewritten as follows:

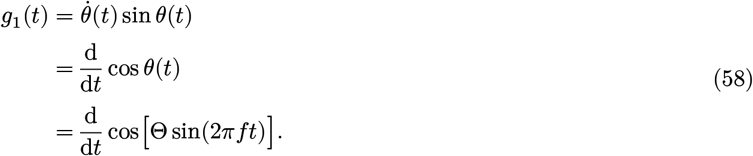

Next, using the Jacobi-Anger expansion [4] the composition cos[Θ sin(2*πft*)] can be rewritten as:

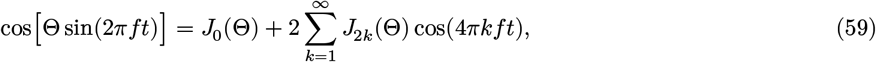

where 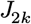 is the Bessel function of the first kind of order 2*k*. Substituting this into Equation 58 the function *g*_1_(*t*) becomes:

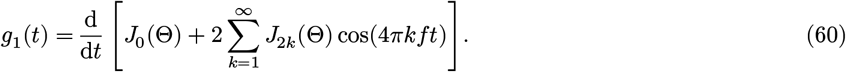

Finally, differentiating the summation term-by-term then yields:

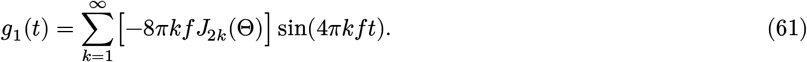

The Fourier coefficients *a*_*n*_ and *b*_*n*_ can then be found using Equations 56 and 61, by comparing the sin(2*πnft*) and cos(2*πnft*) terms, which yields:

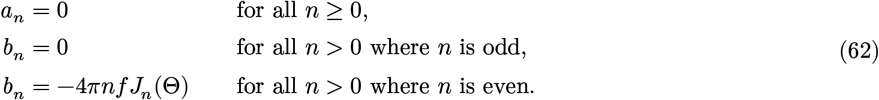

Thus, only the even harmonics (*n* = 2, 4, 6, …) of the sine (odd) components of the expansion of *g*_1_(*t*) are non-zero.

Similar to the expression for *g*_1_(*t*), by applying chain rule and using the fact that *θ*(*t*) = Θ sin(2*πft*) the expression for *g*_2_(*t*) can be rewritten as:

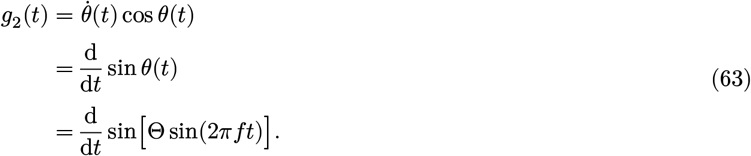

Next, using the Jacobi-Anger expansion [4] the composition sin[Θ sin(2*πft*)] can be rewritten as:

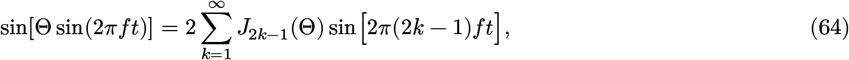

where 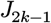 is the Bessel function of the first kind of order 2*k* − 1. After substituting this into Equation 58 the function *g*_2_ becomes:

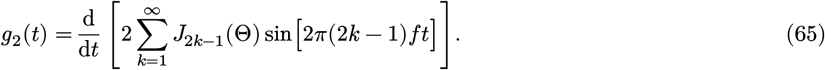

Finally, differentiating the summation term-by-term then yields:

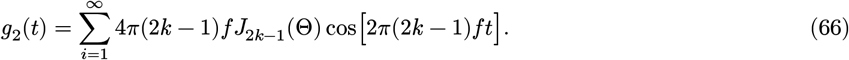

The Fourier coefficients *a*_*n*_ and *b*_*n*_ can then be found using Equations 56 and 66, by comparing the sin(2*πnft*) and cos(2*πnft*) terms, which yields:

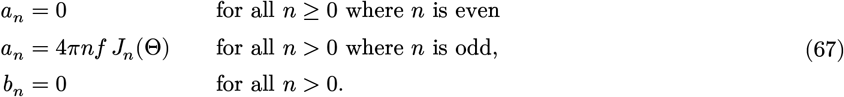

In this case, only the odd harmonics (*n* = 1, 2, 3, …) of the cosine (even) components of the expansion of *g*_2_(*t*) are non-zero.

Finally, when *n* is an integer, the Bessel functions of the first kind [4] are given by:

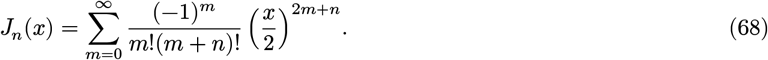

Considering the leading powers of *x* gives:

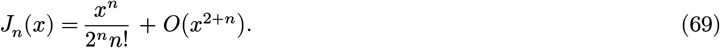

Thus, in for the case *n* = 1;

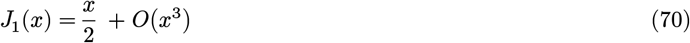

and in the case *n* = 2:

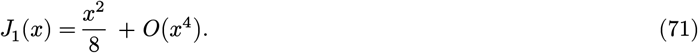

## Notes

### Competing Interest Statement

The authors have declared no competing interest.

## References

1. Pringle, J. W. S. The gyroscopic mechanism of the halteres of Diptera. Phil. Trans. R. Soc. Lond. B 233, 347–384 (1948).

2. Pringle, J. W. S. Insect Flight. (Cambridge University Press, 1957).

3. Nalbach, G. The halteres of the blowfly Calliphora: I. Kinematics and dynamics. J. of Comp. Physiol. A 173, 293–300 (1993).

4. Grimaldi, D., Engel, M. S., Engel, M. S. & Engel, S. C. and P. M. S. Evolution of the Insects. (Cambridge University Press, 2005).

5. Marshall, S. A. Flies: The Natural History & Diversity of Diptera. (Firefly Books, 2012).

6. McAlister, E. The Secret Life of Flies. (Natural History Museum, London, 2017).

7. Fraenkel, G. & Pringle, J. W. S. Biological Sciences: Halteres of flies as gyroscopic organs of equilibrium. Nature 141, 919–920 (1938).

8. Agrawal, S., Grimaldi, D. & Fox, J. L. Haltere morphology and campaniform sensilla arrangement across Diptera. Arth. Struc. & Develop. 46, 215–229 (2017).

9. Gnatzy, W., Grünert, U. & Bender, M. Campaniform sensilla of Calliphora vicina (Insecta, Diptera). Zoomorph. 106, 312–319 (1987).

10. Smith, D. S. The fine structure of haltere sensilla in the blowfly Calliphora erythrocephala (Meig.), with scanning electron microscopic observations on the haltere surface. Tissue and Cell 1, 443–484 (1969).

11. Dickerson, B. H., de Souza, A. M., Huda, A. & Dickinson, M. H. Flies regulate wing motion via active control of a dual-function gyroscope. Cur. Biol. 29, 3517–3524.e3 (2019).

12. Heide, G. Neural mechanisms of flight control in Diptera. BIONA-report 2, 35–52 (1983).

13. Tu, M. S. & Dickinson, M. H. Modulation of negative work output from a steering muscle of the blowfly Calliphora vicina. J. Exp. Biol. 192, 207–224 (1994).

14. Nalbach, G. & Hengstenberg, R. The halteres of the blowfly Calliphora: II. Three-dimensional organization of compensatory reactions to real and simulated rotations. J Comp Physiol A 175, (1994).

15. Dickinson, M. H. Haltere–mediated equilibrium reflexes of the fruit fly, Drosophila melanogaster. Phil. Trans. R. Soc. Lond. B 354, 903–916 (1999).

16. Maimon, G. & Abbott, L. F. Functional logic of a cognitive brain system for navigation. Ann. Rev. Neurosci. 49, 411–434 (2026).

17. Dorkenwald, S. et al. FlyWire: online community for whole-brain connectomics. Nat. Methods 19, 119–128 (2022).

18. Schlegel, P. et al. Whole-brain annotation and multi-connectome cell typing of Drosophila. Nature 634, 139–152 (2024).

19. Götz, K. G. Course-control, metabolism and wing interference during ultralong tethered flight in Drosophila melanogaster. J. Exp. Biol. 128, 35–46 (1987).

20. Ehrhardt, E. et al. Single-cell type analysis of wing premotor circuits in the ventral nerve cord of Drosophila melanogaster. Preprint at 10.1101/2023.05.31.542897 (2023).

21. Lehmann, F. O. & Dickinson, M. H. The changes in power requirements and muscle efficiency during elevated force production in the fruit fly Drosophila melanogaster. J. Exp. Biol. 200, 1133–1143 (1997).

22. Muijres, F. T., Iwasaki, N. A., Elzinga, M. J., Melis, J. M. & Dickinson, M. H. Flies compensate for unilateral wing damage through modular adjustments of wing and body kinematics. Interface Focus 7, 20160103 (2017).

23. Sherman, A. & Dickinson, M. H. A comparison of visual and haltere-mediated equilibrium reflexes in the fruit fly Drosophila melanogaster. J. Exp. Biol. 206, 295–302 (2003).

24. Nalbach, G. Extremely non-orthogonal axes in a sense organ for rotation: Behavioural analysis of the dipteran haltere system. Neurosci. 61, 149–163 (1994).

25. Sherman, A. & Dickinson, M. H. Summation of visual and mechanosensory feedback in Drosophila flight control. J. Exp. Biol. 207, 133–142 (2004).

26. Ristroph, L. et al. Discovering the flight autostabilizer of fruit flies by inducing aerial stumbles. PNAS 107, 4820–4824 (2010).

27. Dickinson, M. H. & Muijres, F. T. The aerodynamics and control of free flight manoeuvres in Drosophila. Phil. Trans. R. Soc. B 371, 20150388 (2016).

28. Beatus, T., Guckenheimer, J. M. & Cohen, I. Controlling roll perturbations in fruit flies. J. R. Soc. Interface. 12, 20150075 (2015).

29. Whitehead, S. C. et al. Neuromuscular embodiment of feedback control elements in Drosophila flight. Sci. Adv. 8, eabo7461 (2022).

30. Whitehead, S. C., Beatus, T., Canale, L. & Cohen, I. Pitch perfect: how fruit flies control their body pitch angle. J. Exp. Biol. 218, 3508–3519 (2015).

31. Giraldo, Y. M. et al. Sun navigation requires compass neurons in Drosophila. Curr. Biol. 28, 2845–2852.e4 (2018).

32. Weir, P. T. & Dickinson, M. H. Functional divisions for visual processing in the central brain of flying Drosophila. PNAS 112, E5523–E5532 (2015).

33. Hulse, B. K., Stanoev, A., Turner-Evans, D. B., Seelig, J. D. & Jayaraman, V. A rotational velocity estimate constructed through visuomotor competition updates the fly’s neural compass. 2023.09.25.559373 Preprint at 10.1101/2023.09.25.559373 (2023).

34. Hulse, B. K. et al. A connectome of the Drosophila central complex reveals network motifs suitable for flexible navigation and context-dependent action selection. eLife 10, e66039 (2021).

35. Turner-Evans, D. et al. Angular velocity integration in a fly heading circuit. eLife 6, e23496 (2017).

36. Green, J. et al. A neural circuit architecture for angular integration in Drosophila. Nature 546, 101–106 (2017).

37. Muijres, F. T., Elzinga, M. J., Melis, J. M. & Dickinson, M. H. Flies evade looming targets by executing rapid visually directed banked turns. Science 344, 172–177 (2014).

38. Kathman, N. D. & Fox, J. L. Representation of haltereoscillations and integration with visual Inputs in the fly central complex. J. Neurosci. 39, 4100–4112 (2019).

39. David, C. T. Visual control of the partition of flight force between lift and thrust in free-flying Drosophila. Nature 313, 48–50 (1985).

40. David, C. T. The relationship between body angle and flight speed in free-flying Drosophila. Physiol. Entomol. 3, 191–195 (1978).

41. Schilstra, C. & van Hateren, J. H. Stabilizing gaze in flying blowflies. Nature 395, 654–654 (1998).

42. Hengstenberg, R. Gaze control in the blowfly Calliphora: a multisensory, two-stage integration process. Sem. Neurosci. 3, 19–29 (1991).

43. Tammero, L. F. & Dickinson, M. H. The influence of visual landscape on the free flight behavior of the fruit fly Drosophila melanogaster. J. Exp. Biol. 205, 327–343 (2002).

44. Censi, A., Straw, A. D., Sayaman, R. W., Murray, R. M. & Dickinson, M. H. Discriminating external and internal causes for heading changes in freely flying Drosophila. PLOS Comp. Biol. 9, e1002891 (2013).

45. Muijres, F. T., Elzinga, M. J., Iwasaki, N. A. & Dickinson, M. H. Body saccades of Drosophila consist of stereotyped banked turns. J. Exp. Biol. 218, 864–875 (2015).

46. van Breugel, F., Jewell, R. & Houle, J. Active anemosensing hypothesis: how flying insects could estimate ambient wind direction through sensory integration and active movement. J. R. Soc. Inter. 19, 20220258.

47. May, C. E., Cellini, B., van Breugel, F. & Nagel, K. I. A compact multisensory representation of self-motion is sufficient for computing an external world variable. Preprint at 10.1101/2025.05.09.653128 (2025).

48. Stupski, S. D. & Breugel, F. van. Wind gates olfaction-driven search states in free flight. Cur. Biol. 34, 4397–4411.e6 (2024).

49. Cellini, B., Boyacıoğlu, B., Stupski, S. D. & Breugel, F. van. Discovering and exploiting active sensing motifs for estimation with empirical observability. 2024.11.04.621976 Preprint at 10.1101/2024.11.04.621976 (2024).

50. Seelig, J. D. & Jayaraman, V. Neural dynamics for landmark orientation and angular path integration. Nature 521, 186–191 (2015).

51. Kutschireiter, A., Basnak, M. A., Wilson, R. I. & Drugowitsch, J. Bayesian inference in ring attractor networks. PNAS 120, e2210622120 (2023).

52. Tu, M. S. & Dickinson, M. H. The control of wing kinematics by two steering muscles of the blowfly (iCalliphora vicina). J. Comp. Physiol. A 178, 813–830 (1996).

53. Balint, C. N. & Dickinson, M. H. Neuromuscular control of aerodynamic forces and moments in the blowfly, Calliphora vicina. J. Exp. Biol. 207, 3813–3838 (2004).

54. Seelig, J. D. & Jayaraman, V. Feature detection and orientation tuning in the Drosophila central complex. Nature 503, 262–266 (2013).

55. Fisher, Y. E. Flexible navigational computations in the Drosophila central complex. Curr. Opin. Neurobiol. 73, 102514 (2022).

56. Fisher, Y. E., Marquis, M., D’Alessandro, I. & Wilson, R. I. Dopamine promotes head direction plasticity during orienting movements. Nature 612, 316–322 (2022).

57. Kim, S. S., Hermundstad, A. M., Romani, S., Abbott, L. F. & Jayaraman, V. Generation of stable heading representations in diverse visual scenes. Nature 576, 126–131 (2019).

58. Fisher, Y. E., Lu, J., D’Alessandro, I. & Wilson, R. I. Sensorimotor experience remaps visual input to a heading-direction network. Nature 576, 121–125 (2019).

59. Kim, S. S., Rouault, H., Druckmann, S. & Jayaraman, V. Ring attractor dynamics in the Drosophila central brain. Science 356, 849–853 (2017).

60. Green, J., Vijayan, V., Mussells Pires, P., Adachi, A. & Maimon, G. A neural heading estimate is compared with an internal goal to guide oriented navigation. Nat. Neurosci. 22, 1460–1468 (2019).

61. Hughes, T. P. Elmer Sperry. (Johns Hopkins University Press, 1971).

62. David, C. T. Compensation for height in the control of groundspeed byDrosophila in a new, ‘barber’s pole’ wind tunnel. J. Comp. Physiol. 147, 485–493 (1982).

63. Suver, M. P. et al. Encoding of wind direction by central neurons in Drosophila. Neuron 102, 828–842.e7 (2019).

64. Matheson, A. M. M. et al. A neural circuit for wind-guided olfactory navigation. Nat. Comm. 13, 4613 (2022).

65. Honkanen, A., Adden, A., da Silva Freitas, J. & Heinze, S. The insect central complex and the neural basis of navigational strategies. J. Exp. Biol. 222, (2019).

66. van Breugel, F., Jewell, R. & Houle, J. Active anemosensing hypothesis: how flying insects could estimate ambient wind direction through sensory integration and active movement. J. R. Soc. Inter. 19, 20220258 (2022).

67. Lyu, C., Abbott, L. F. & Maimon, G. Building an allocentric travelling direction signal via vector computation. Nature 601, 92–97 (2022).

68. Stone, T. et al. An anatomically constrained model for path integration in the bee brain. Current Biology 27, 3069–3085 (2017).

69. Baker, K. L. et al. Algorithms for olfactory oearch across species. J. Neurosci. 38, 9383–9389 (2018).

70. Dana, H. et al. High-performance calcium sensors for imaging activity in neuronal populations and microcompartments. Nat. Meth. 16, 649–657 (2019).

71. Jenett, A. et al. A GAL4-driver line resource for Drosophila neurobiology. Cell reports 2, 991–1001 (2012).

72. Diao, F. et al. Plug-and-play genetic access to Drosophila cell types using exchangeable exon cassettes. Cell Reports 10, 1410–1421 (2015).

73. Dickinson, M. H.Lehmann, F.-O. & Gotz, K. G. The active control of wing rotation by Drosophila. J. Exp. Biol. 182, 173–189 (1993).

74. Ros, I. G., Omoto, J. J. & Dickinson, M. H. Descending control and regulation of spontaneous flight turns in Drosophila. Curr. Biol. 34, 531–540.e5 (2024).

75. Suver, M. P., Mamiya, A. & Dickinson, M. H. Octopamine neurons mediate flight-induced modulation of visual processing in Drosophila. Curr. Biol. 22, 2294–2302 (2012).

76. Wolff, T., Iyer, N. A. & Rubin, G. M. Neuroarchitecture and neuroanatomy of the Drosophila central complex: A GAL4-based dissection of protocerebral bridge neurons and circuits. J. Comp. Neurol. 523, 997–1037 (2015).

77. Reiser, M. B. & Dickinson, M. H. A modular display system for insect behavioral neuroscience. J. Neurosci. Meth. 167, 127–139 (2008).

## Supplementary references

[1] Nalbach, G., and Hengstenberg, R., 1993, “The Halteres of the Blowfly calliphora, I. Kinematics and Dynamics.,” J. Comp. Physiol A, 173, pp. 293–300. 10.1007/BF00191842.

[2] Pringle, J. S., 1948, “The Gyroscopic Mechanism of the Halteres of Diptera.,” Philosophical Transactions of the Royal Society of London. Series B, Biological Sciences, 233, p. 347––384. 10.1007/BF00191842.

[3] Vaxenburg, R., Siwanowicz, I., Merel, J., Robie, A., Morrow, C., Novati, G., Stefanidi, Z., Both, G., Gwyneth M Card, G. M., Reiser, M. B., M Botvinick, M. M., Branson, K. M., Tassa, Y., and Turaga, S. C., 2025, “Whole-Body Physics Simulation of Fruit Fly Locomotion,” Nature, 643, pp. 1312–1320. 10.1038/s41586-025-09029-4.

[4] Abramowitz, M., and Stegun, I. A., 1964, Handbook of Mathematical Functions with Formulas, Graphs, And Mathematical Tables, Dover Publications.

